# Improving the workflow to crack Small, Unbalanced, Noisy, but Genuine (SUNG) datasets in bioacoustics: the case of bonobo calls

**DOI:** 10.1101/2022.06.26.497684

**Authors:** Vincent Arnaud, François Pellegrino, Sumir Keenan, Xavier St-Gelais, Nicolas Mathevon, Florence Levréro, Christophe Coupé

## Abstract

Despite the accumulation of data and studies, deciphering animal vocal communication remains highly challenging. While progress has been made with some species for which we now understand the information exchanged through vocal signals, researchers are still left struggling with sparse recordings composing Small, Unbalanced, Noisy, but Genuine (SUNG) datasets. SUNG datasets offer a valuable but distorted vision of communication systems. Adopting the best practices in their analysis is therefore essential to effectively extract the available information and draw reliable conclusions. Here we show that the most recent advances in machine learning applied to a SUNG dataset succeed in unraveling the complex vocal repertoire of the bonobo, and we propose a workflow that can be effective with other animal species. We implement acoustic parameterization in three feature spaces along with three classification algorithms (Support Vector Machine, xgboost, neural networks) and their combination to explore the structure and variability of bonobo calls, as well as the robustness of the individual signature they encode. We underscore how classification performance is affected by the feature set and identify the most informative features. We highlight the need to address data leakage in the evaluation of classification performance to avoid misleading interpretations. Finally, using a Uniform Manifold Approximation and Projection (UMAP), we show that classifiers generate parsimonious data descriptions which help to understand the clustering of the bonobo acoustic space. Our results lead to identifying several practical approaches that are generalizable to any other animal communication system. To improve the reliability and replicability of vocal communication studies with SUNG datasets, we thus recommend: i) comparing several acoustic parameterizations; ii) adopting Support Vector Machines as the baseline classification approach; iii) explicitly evaluating data leakage and possibly implementing a mitigation strategy; iv) visualizing the dataset with UMAPs applied to classifier predictions rather than to raw acoustic features.

## Introduction

Cracking animal vocal communication codes has always constituted a motivating challenge for bioacousticians, evolutionary biologists and ethologists, and impressive breakthroughs have been achieved in the last decade thanks to advances in both data collection and machine learning, in conjunction with well-designed experimental approaches (e.g. Coye, et al., 2016; Turesson, et al., 2016; Mielke & Zuberbuhler, 2013). Building upon these success stories, can we conjecture that the puzzle of animal vocal communications will be soon solved? Probably not really. The high level of understanding achieved for a few dozens of species should not obscure the fact that for the vast majority of animal species, not much is known yet. For most animal species, neat, clean, and massive datasets are out of reach and bioacousticians have to cope with Small, Unbalanced, Noisy, but Genuine (SUNG) datasets characterized by data paucity, unbalanced number of recordings across individuals, contexts, or call categories for instance, and by noisy and sometimes reverberant recording environments. Despite their imperfection, the worth of such datasets, which are often expensive to collect and annotate, is invaluable as they inform us about the complex communication systems of animal species that may be dramatically endangered or of major scientific interest. As a result, it is becoming increasingly important to adopt the best possible practices in the analysis of such datasets, both to provide reproducible studies and to reach robust and reliable conclusions about the species communication systems, and beyond that, about more general questions concerning the evolution, diversity, and complexity of communication systems.

In mammals, individuals often produce vocalizations potentially informative to their conspecifics in their “here and now” context of emission. Additionally, these signals can also provide idiosyncratic clues to the emitter identity, which is often an essential information for territorial defense, social interaction and cohesion (e.g. Charlton, et al., 2020). In social species, especially those living in fission-fusion systems (i.e. the size and composition of the social group change as time passes and animals move throughout the environment; animals merge into a group (fusion) or split (fission)), the “who” is therefore as important as the “what” in vocal communication. Much research has therefore sought to determine which acoustic primitives (a.k.a. features) encode the “who” and the “what” respectively in order to test hypotheses about the functions fulfilled by vocal communication, through playback experiments, and more recently resynthesis for instance (Charlton, et al., 2012). While straightforward communication systems where the emitter identity and the contextual information are sequentially encoded (a strategy called temporal segregation) are attested (Jansen, Cant, & Manser, 2012), the acoustic features on which the context-specific information and the vocal signature develop differ across species, and their identification can be much more complex and challenging (Clay, Archbold, & Zuberbühler, 2015; Fischer, Wadewitz, & Hammerschmidt, 2017).

A key step to disentangle the “who” from the “what” is thus to assess the discriminative power of potential acoustic features by automatic classification in order to infer their putative role in communication as signals, carrying some information about the identity or intent of the emitter, the call type, and the context of utterance. Fifteen years ago, Roger Mundry and Christina Sommer introduced permutated Discriminant Function Analysis (pDFA) as a means of assessing the statistical power of a set of acoustic properties in a discrimination task (Mundry & Sommer, 2007). This method belongs to the field of supervised machine learning, where a classifier is trained to discriminate data samples (training set) according to a priori categories (labels). The performance of the classifier is then measured in terms of its ability to correctly generalize the discrimination decision to new unseen samples belonging to the same categories (test set). A common – but sometimes overlooked – detrimental issue occurs when the classifier’s decision is correct, but based on faulty premises because the samples in the training and test sets are not drawn from independent datasets and share confounding properties other than the category label itself (e.g., a background acoustic noise that leaks information about the identity of the recorded individual), a phenomenon known as data leakage in data mining and machine learning (Kaufman, Rosset, & Perlich, 2012; see also the “Husky vs Wolf’’ experiment in Ribeiro, Singh & Guestrin, 2016, for a striking illustration). The pDFA was specifically designed to mitigate the risk of overestimating discriminability due to such a non-independence artifact, although it is not immune to such shortcomings. It has become the standard approach for bioacoustic analysis and it is still routinely used nowadays despite its limitations. Specifically, it is neither the best nor the most accurate classification algorithm for assessing discriminability with SUNG datasets. As a paradoxical consequence, pDFA can be expected, on the one hand, to underestimate the information present in a dataset (because of suboptimal classification), and, on the other hand, to overestimate the class discriminability (because of residual non-independence). Meanwhile, in other scientific fields, impressive improvements have been made to address similar issues, by implementing more powerful statistical and machine-learning algorithms in more controlled configurations of evaluation. As aforementioned, such algorithms, including deep-learning neural networks, have recently been successfully applied to animal communication, mainly in situations where data paucity was not an issue (e.g., Aodha, et al., 2018, but see Turesson, et al., 2016 and Clink & Klinck, 2019 for applications to smaller datasets; Stowell, 2022 for a recent review; Valletta, et al., 2017 for a broader perspective).

In addition to automatic classification, graphical exploration of a corpus of animal calls projected into an informative feature space is often a helpful – and essential – step in understanding the structure of their repertoire. Since multiple acoustic features are usually involved, a reduction in dimensionality is required to get a human-friendly low-dimensional representation, usually in two (plane) or three (volume) dimensions. While this reduction was traditionally achieved through linear or near-linear transformations, such as Principal Component Analysis or Multidimensional Scaling, innovative non- linear approaches such as t-distributed stochastic neighborhood embedding (t-SNE, Van der Maaten & Hinton, 2008) and Uniform Manifold Approximation and Projection (UMAP, McInnes, Healy, & Melville, 2020) have recently emerged. These methods generally result in intuitive representations of the local structure present in complex datasets, at the expense of the significance of the global structure. Both t- SNE and UMAP have already been successfully applied to animal communication, either as exploratory methods to assess repertoire discreteness vs. grading, or to compare the relevance of several feature sets (see Goffinet, et al., 2021; Smith-Vidaurre, Araya-Salas, & Wright, 2020; Valente, et al., 2019; Valente, et al., 2022 for examples; Sainburg, Thielk, & Gentner, 2020 and Thomas, et al., 2022 for thorough discussions of the potential benefits of these methods, as well as their limits with small datasets).

In this paper, we apply an automatic classification workflow to a SUNG dataset whose structure should seem fairly conventional to most bioacousticians: several recordings of calls produced in sequences of varying lengths, belonging to half a dozen types, and produced by a dozen individuals. This dataset consists of audio recordings of calls emitted by captive bonobos (*Pan paniscus*). It provides a case study where the identification of individuals on the basis of their vocalizations and the identification of call types is not trivial. The bonobo vocal repertoire was described several decades ago in two seminal studies that highlighted its graded nature (Bermejo & Omedes, 1999; de Waal, 1988). It is structured on almost a dozen prototypical types that conjugate modulated voiced vocalizations with scream components, also exhibiting nonlinear phenomena. Although quantitative studies with bonobos are still rare, it has recently been shown that an individual signature is detectable in bonobo vocalizations and that the reliability of this signature differs from one call type to another (Keenan, et al., 2020). Here we implement a systematic comparison of several classification approaches to assess the strength and stability of individual bonobo signatures in the vocal repertoire. Our results establish whether the level of performance is the result of a lack of intrinsically encoded information in the vocalizations or a suboptimal classification approach.

The research question we have addressed is therefore: can state-of-the-art automatic classification approaches lead to a more accurate estimation of the information encoded in a SUNG dataset and to a more comprehensive understanding of the functioning of an animal communication system? We present and evaluate the relevance of several methods that can be used to overcome the difficulties inherent in raw naturalistic dataset. In the end, the objective of this paper is to propose an operational workflow that can be used by bioacousticians in concrete situations.

Figure 1 gives a graphical overview of the proposed methodology and the paper is organized as follows:

**Figure 1.**
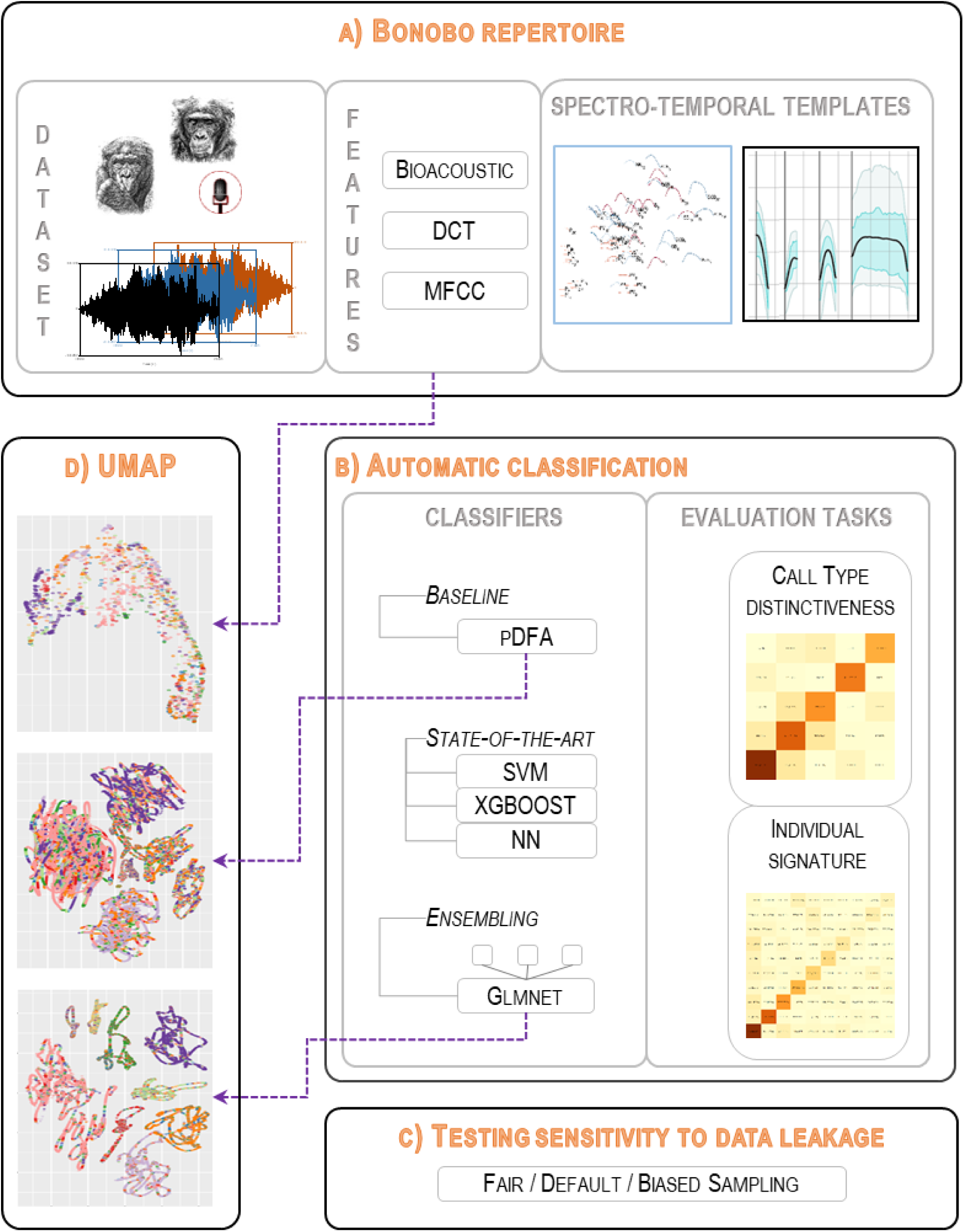
Workflow implemented to analyze a dataset of animal vocalizations. A. Three different sets of acoustic features are extracted from the dataset by different methods (BIOACOUSTIC, DCT and MFCC) and then used to deduce call type templates. B. The performance of three state-of-the-art classifiers and their ensembling combinations is assessed and compared to that of a discriminant analysis (pDFA) in two tasks: identification of call types (bonobos have a vocal repertoire composed of different calls) and discrimination between emitters (identification of individual vocal signatures). C. The accuracy sensitivity to the composition of the training and test sets and to the induced data leakage is then evaluated. D. Finally, we discuss the use of classifier outputs to improve low-dimensional representations of the call acoustic space (UMAPs).

In section I, we present the repertoire of bonobo calls and the SUNG nature of our audio dataset. We define three sets of acoustic parameters to improve the robustness of the analysis of noisy vocalizations.

In section II, we test three classification algorithms that are compatible with the limited amount of data available and its imbalance between categories, and we carefully choose adequate performance metrics. We also build combined/stacked classifiers to test the complementarity between parameter sets and classification algorithms, with the aim of improving the overall robustness of the analysis compared to a baseline obtained by a pDFA approach classically used in bioacoustics (Mundry & Sommer, 2007).

In section III, we provide an in-depth evaluation of the impact of data leakage due to non-independence on classification results. We demonstrate why data leakage should not be overlooked by reporting the performances reached when maximizing the independence between the training and test datasets, compared to results from random partitions or from a maximally dependent (purposefully ill-designed) partition. We show how the use of a genetic algorithm allows the construction of training and test datasets leading to an almost unbiased accuracy estimation.

Section IV illustrates the visual improvement in terms of UMAP representation when an appropriate classifier is applied as a preprocessor to get informative latent features over the representation directly extracted from the acoustic parameters.

Section V discusses the main achievements and limitations of our study.

In section VI, we detail the methodology and discuss how bioacousticians can apply this data science approach to explore biological questions.

Datasets and analysis codes are made available on http://github.com/keruiduo/SupplMatBonobos

### I. The bonobo recordings: a SUNG dataset

#### A quick overview of bonobo vocal communication

Vocal communication is ubiquitous in bonobos and different call types are used in a flexible way across contexts and activities, resulting in complex and meaningful combinations (e.g. inter-party travel recruitment; food preference; see de Waal, 1988, Clay & Zuberbühler, 2009; Schamberg, et al., 2017, among others). The bonobo vocal repertoire is complex and graded: acoustic characteristics can vary within each call type, and the different types are distributed along an acoustic continuum. Two descriptions of call types have been proposed and are largely converging (Bermejo & Omedes, 1999; de Waal, 1988). Most calls are confidently labeled by experts as belonging to one of 12 types (the ten shown in Figure 2, plus grunts and screams), despite the gradual changes that lead to some degree of uncertainty. However, doing this automatically is a much more difficult challenge than in other primate species with a discrete repertoire (e.g. common marmosets, Oikarinen, et al., 2019; Turesson, et al., 2016). This classification is based on the characteristics of the tonal contour when detectable (general shape and duration) and the vocal effort (illustrated by the presence of energy in the higher harmonics and nonlinear phenomena).

**Figure 2.**
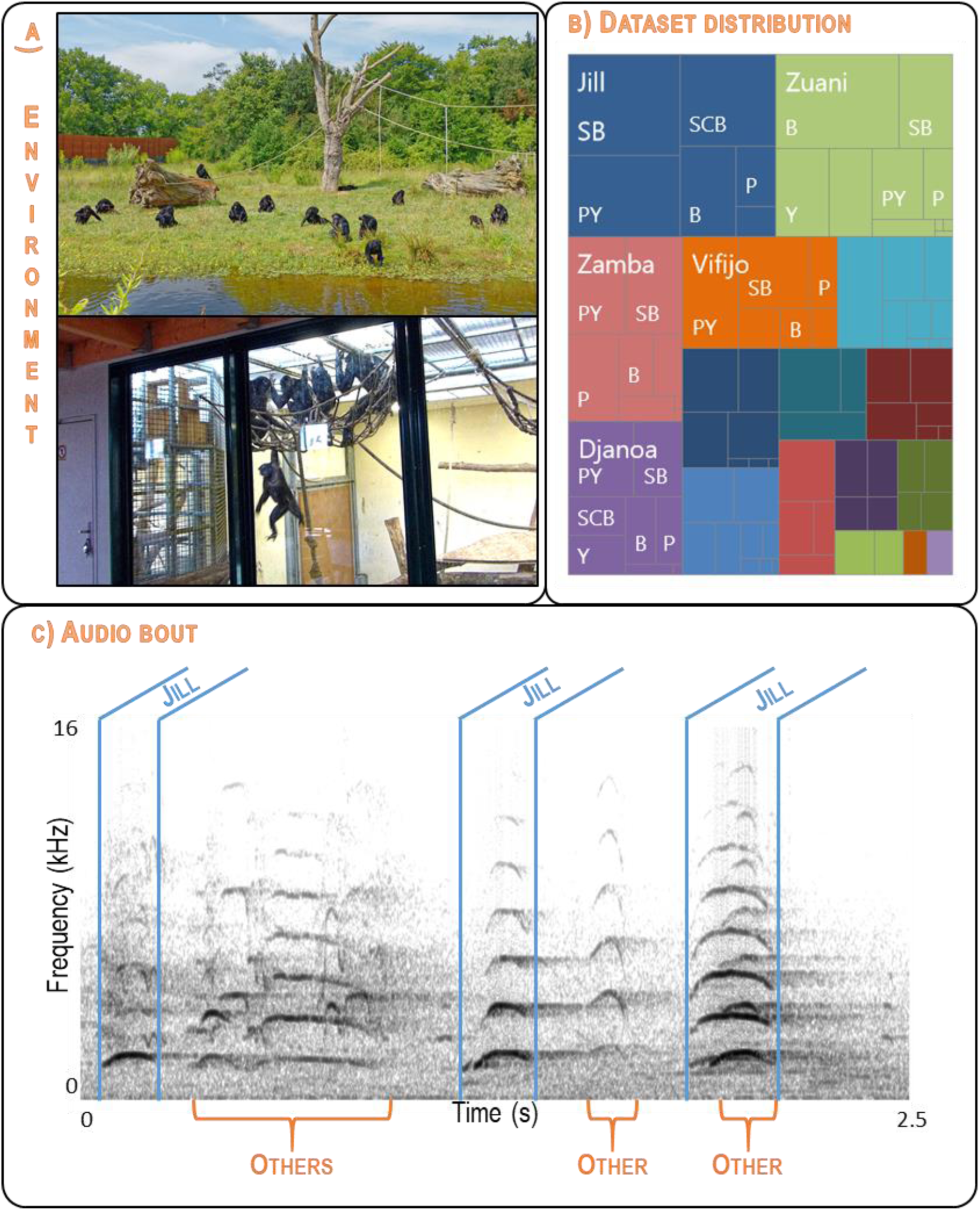
An example of a SUNG bioacoustic dataset: recordings of bonobo calls in social contexts. A. Some individuals are recorded in outdoor enclosures while others are recorded inside buildings. B. The number of calls varies between individuals (unbalanced distribution coded by colored rectangles) and by call type (coded by internal rectangles for each individual). The five most-represented individuals are named. The four least represented individuals are not shown on the chart. C. Spectrogram of a typical recorded bout (2.5 seconds extracted from the Jill698 recording) showing the difficulty of isolating good quality calls. A sequence of three calls produced by Jill can be identified (sections delimited by blue boundaries). Other individuals vocalize in the background (sections marked with orange curly brackets). Jill’s third call is not analyzed as it overlaps too much with other vocalizations. Photo credits: F. Levrero (Top) & F. Pellegrino (Bottom).

Besides the “what” contextual information, the “who” information is crucial to navigate the complex fission-fusion society of bonobos. Recent research suggests, however, that the individual vocal signature is more salient in high-arousal calls than in low-arousal calls (Keenan, et al., 2020). According to this result, the identification of an individual would therefore be easier on the basis of high-arousal calls. However, playback experiments have shown that bonobos are able to discriminate a familiar from an unfamiliar congener based on low-arousal calls (peep-yelp; Keenan, et al., 2016). Note that this kind of individual signature depending on the type of call had already been reported in other mammal species (Bouchet, et al., 2012; Leliveld, Scheumann, & Zimmermann, 2011).

A SUNG dataset is characterized by data paucity, the imbalance in the number of recordings between individuals, between contexts or between call types, and by noisy and sometimes reverberant recording environments. A first set of these constraints is inherent in the field work conditions, and is not specific to bonobos. Whether in zoos or in the wild, recordings of vocalizations often involve few individuals (typically less than two dozen), in an unbalanced proportion of call types and individuals. Large corpus such as that described in Agamaite, et al. (2015), with dozens of thousands of calls, are therefore very rare. In addition, recordings are always performed in environments with unique characteristics in terms of background soundscape and noise. All these aspects have consequences on the acoustic feature extraction and on the automatic classification, by limiting the amount and quality of data available and exposing the evaluation to potential misinterpretation due to data leakage. These obstacles are well identified in ecology and ethology studies (see for instance Stowell, et al., 2019, for an elegant proposal on acoustic feature extraction and modeling in the context of bird identification), but they remain problematic, in stark contrast to studies on human language for which massive data are now available (see Filippidou & Moussiades, 2020 for a recent comparison of automatic speech recognition systems, for instance).

Further limitations arise from the fact that, in primate graded vocal communication, the functional role of the signals contributing to the vocal repertoire is only partially understood, leading to a potential lack of ground truth (or gold standard) against which automatic classification approaches can be judged. This is definitely true when classifying a bonobo call as belonging to a given type within the species’ graded repertoire, as opposed to an ungraded or less-graded repertoire (e.g., Turesson, et al., 2016). This gold standard problem disappears when automatically identifying the call emitter, whose correct identity is known when the action has been directly observed in absence of overlaps between emitters. But even this supposedly easy situation can be complex in bonobos, as their vocal activity is often unpredictable, making the detection of the emitter difficult among all group members. In addition, a first emitter often triggers vocalizations from a few other individuals, resulting in a sudden intense vocal activity (with overlapping vocalizations). In such situations, unambiguous assignment of an emitter identity to a call produced in a sequence is sometimes impossible, and it requires a lot of recording time to obtain enough calls for which the emitter identity is unambiguously known. Moreover, even if the emitter’s identity is known, deciding whether an acoustic feature is relevant or not is not straightforward. To illustrate, let’s imagine that the acoustic feature A allows an automatic classifier to perfectly identify one individual when it is the emitter, but poorly performs in recognizing all other individuals. On the contrary, considering only feature B leads to a slightly better-than-chance identification for all individuals. Which feature is the most important from the animal’s point of view? In this case, the answer is probably both, but this example aims to highlight that the choice of the right evaluation metrics (average recognition rate, accuracy, etc.) among the dozens available may influence the resulting ethological interpretation (see Methodology subsection “Performance Metrics”).

In this study, we use a dataset from the corpus analyzed in Keenan et al. (2020). It comes from 20 adult bonobos housed in three zoos (Apenheul in The Netherlands, La Vallée des Singes in France and Planckendael in Belgium), totaling 380 hours of recording. Recordings were made during daytime using a Sennheiser MKH70-1 ultradirectional microphone and a Zoom H4 Digital Multitrack Recorder (44.1 kHz sample rate, 16 bits per sample, .wav files), in various contexts (foraging, grooming, aggression, etc. See Keenan, et al., 2020 for details). Vocalizations were manually segmented, identified and then double-checked by two other experimenters with Praat (Boersma, 2006), based on the visual inspection of signals on spectrograms.

The audio quality of the recordings was variable, with many calls being recorded in a reverberant and challenging environment for automatic F0 detection, often leading to uncertainties in Praat. The temporal modulation of F0 was thus derived semi-automatically from the narrow-band spectrograms, thanks to a homemade Praat script based on mouse input of at least two points on the F0 trace on the spectrogram by the experimenter, allowing an interpolated trajectory to be estimated.

Figure 2 illustrates why this dataset can be qualified as SUNG: the recorded environment may be a distant free-ranging enclosure or an indoor room; there is a quite noticeable imbalance in the number of calls per individual, and the audio bouts are noisy, reverberant, and complex to analyze.

We worked with a dataset consisting of 1,971 calls from 20 subjects to perform the preliminary quantitative study to characterize the tonal contour of each type (Section II and Figure 2) (after removing grunts and screams that do not have any tonal component). The following 10 call types are thus described: Peep (P), Yelp (Y), Hiccup (H), Peep Yelp (PY), Soft Bark (SB), Bark (B), Scream Bark (SCB), Whining Whistle (WW), Whistle (W), and Scream Whistle (SCW). For the automatic classification tasks reported in Sections II, III, and IV, we selected the ten individuals for whom at least 70 calls were available from the five most frequent call types Bark (B), Soft Bark (SB), Peep (P), Peep Yelp (PY) and Scream Bark (SCB) (1,560 calls, split by call types and individuals in Table 1).

**Table 1.** Number of calls per individual and per call type in the dataset used for automatic classification. The five call types are: Bark (B), Peep (P), Peep Yelp (PY), Soft Bark (SB), and Scream Bark (SCB).

| Types | Individuals |  |  |  |  |  |  |  |  |  | Total |
| --- | --- | --- | --- | --- | --- | --- | --- | --- | --- | --- | --- |
|  | Bolombo | Busira | Djanao | Hortense | Jill | Kumbuka | Lina | Vifijo | Zamba | Zuani |  |
| B | 6 | 11 | 20 | 9 | 50 | 23 | 8 | 12 | 21 | 114 | 274 |
| P | 27 | 18 | 18 | 26 | 24 | 35 | 24 | 20 | 43 | 20 | 255 |
| PY | 24 | 34 | 49 | 50 | 89 | 26 | 18 | 61 | 56 | 36 | 443 |
| SB | 17 | 5 | 36 | 8 | 112 | 25 | 23 | 51 | 55 | 50 | 382 |
| SCB | 2 | 3 | 22 | 18 | 87 | 0 | 3 | 16 | 18 | 37 | 206 |
| **Total** | 76 | 71 | 145 | 111 | 362 | 109 | 76 | 160 | 193 | 257 | 1,560 |

#### A sketch of across-individual variability

For each call type, we estimated a template (or prototype) of the fundamental frequency (F0) corresponding to the average F0 trajectory estimated over the whole corpus. F0 contour extraction and smoothing was automatically performed in Praat and manually corrected for gross errors (typically subharmonic detection). These F0 templates are presented in Figure 3 for the ten types of tonal call for which F0 can be extracted. It should be noted that the Hiccup (H), Whining Whistle (WW) and Scream Whistle (SCW) are very rare vocal production and their templates thus were estimated from a small number of samples, which led us not to consider them in the following sections. Considering only the bell curve types (P, PY, SB, B, and SCB), a continuum is visible on the F0 dimension with increasing F0 average value and excursion, except between barks and scream barks, which differ mainly in the absence or presence of a screaming component (like deterministic chaos). This aspect is not captured by the F0 trajectory, but it is suggested by the rather large difference in average harmonicity between the two categories, with SCB’s harmonicity being on average 2.2 dB lower than that of B. SCB thus has the lowest harmonicity among the ten types displayed in its energy distribution, leading to a highly salient perceptual roughness.

**Figure 3.**
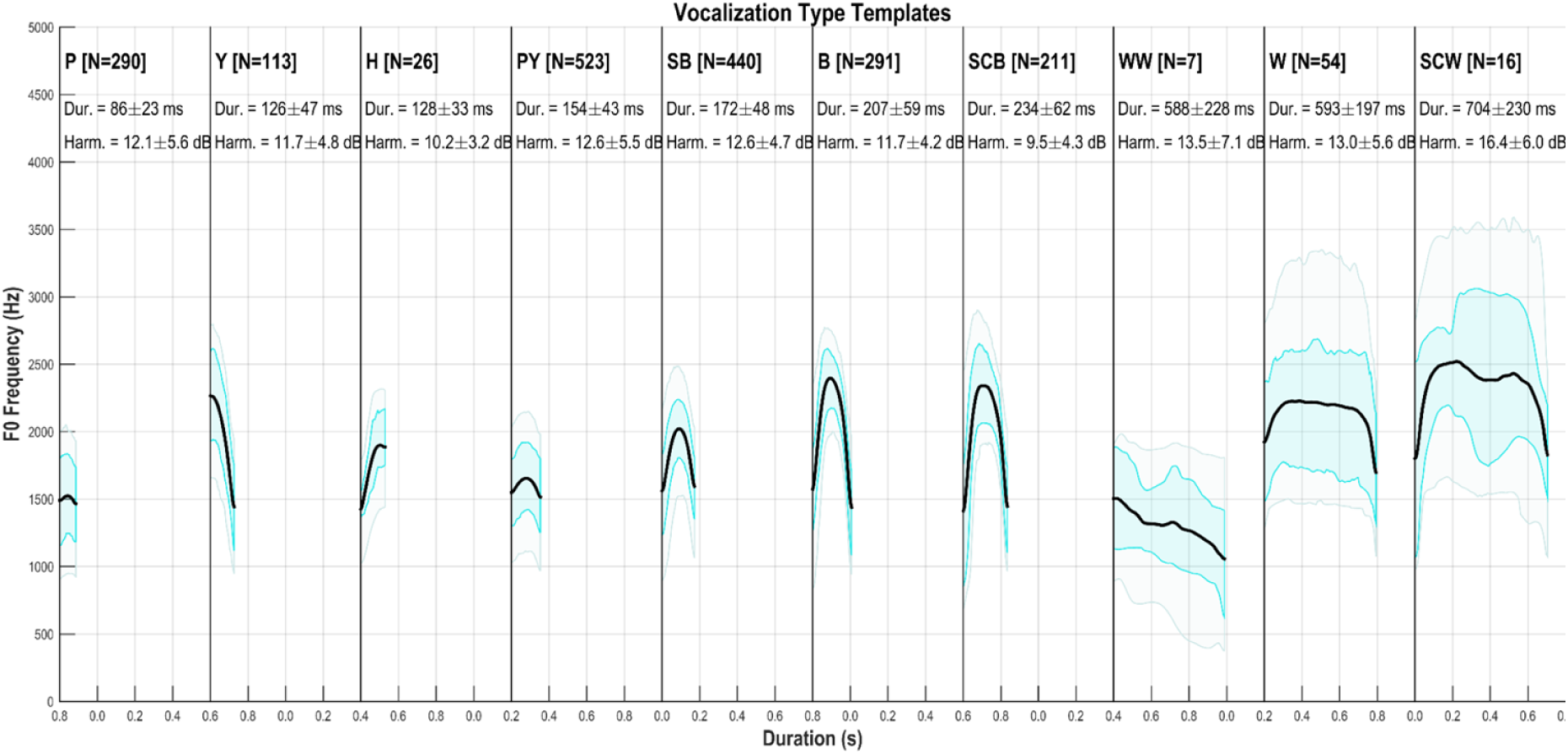
Templates of F0 (pitch) for each call type. The average F0 trajectory (black line) is calculated from all recordings (using PRAAT software). The shaded area covers 50% or 80% of the distribution (blue and grey areas respectively). For each type of call, individual calls were time-scaled to the average duration of the type. N = number of calls analyzed; Dur = call duration (mean and standard deviation, in ms); Harm = harmonicity (mean and standard deviation, in dB). The types are ranked by increasing average duration: Peep (P), Yelp (Y), Hiccup (H), Peep Yelp (PY), Soft Bark (SB), Bark (B), Scream Bark (SCB), Whining Whistle (WW), Whistle (W), and Scream Whistle (SCW).

To further illustrate the graded aspect of the bonobo vocal repertoire, we also computed F0 templates at the individual level. Their distribution is shown in Figure 4 (left), with a miniature of each template (at scale 1/10^th^) represented in a two-dimension space: average F0 and average duration. A large variation in F0 is observed among the individual peep templates, and their short duration distinguishes them from the other types. The Bark type spans a large area of the acoustic space, with a large variation in both duration and F0. For a given individual, the relative weight of the temporal and frequential dimensions may differ, as illustrated in Figure 4 (right) for individuals #19 and #20. On average, the calls produced by individual #20 are higher-pitched than those produced by individual #19, but an additional difference is highlighted by the respective position of their SB templates which is very close to B for #19 while it is more central for #20. This observation suggests that the inter-individual variation observed is not entirely constrained by anatomical differences and that each individual’s repertoire is akin to an idiolect in human linguistics. This type of graded repertoire with overlapping categories represents a difficult challenge for automatic classification methods (Wadewitz, et al., 2015).

**Figure 4.**
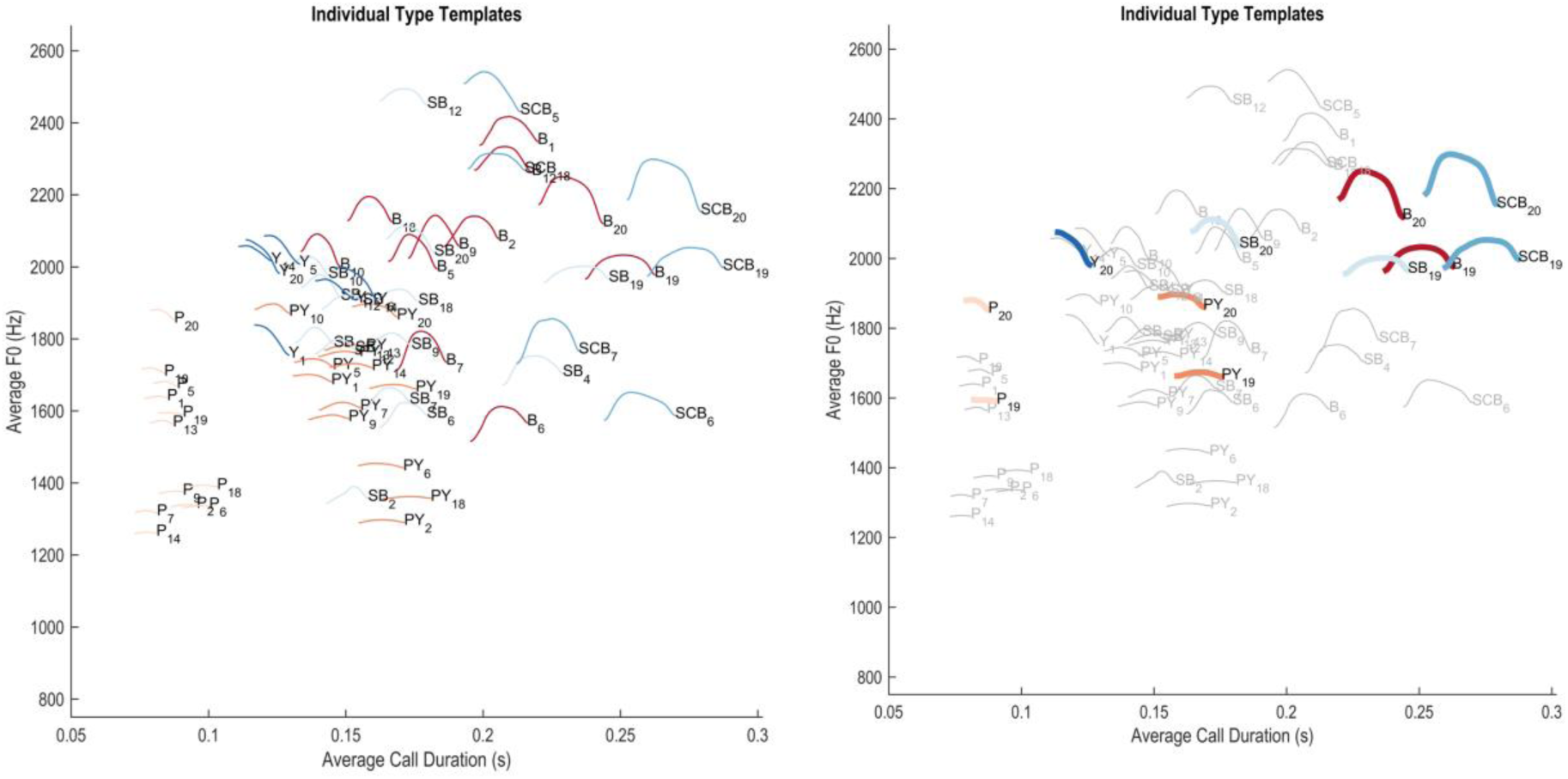
Representation of call F0 templates at the individual level. Each color/hue combination corresponds to a call type (P, Y, PY, SB, B, SCB). Each curve is a 1/10th miniature of an individual’s F0 template. The call type (acronym) and individual identity (numerical index) are indicated. **Left.** All Individuals and call types for which at least 5 samples were available. **Right.** Same figure with the repertoire of individuals #19 and #20 highlighted.

### II. Automatic classification: From pDFA to state-of-the-art approaches

#### Extraction of acoustic features

To parametrize each individual vocalization, we considered three different sets of features: the Bioacoustic set, the MFCC set and the DCT set. They are summarized in the appendix (Table A1). The **Bioacoustic** set adopts a fairly standard approach on primate call analysis, which has already been used in bonobo studies (e.g., Clay & Zuberbühler, 2009). The **MFCC** set, although less common, has already been successfully applied to primate call recognition and individual identification (e.g., Mielke & Zuberbühler, 2013; Spillmann, et al., 2017). This full-spectrum approach may potentially be able to highlight fine-grained spectral differences that are not captured with the standard bioacoustic approach. To our knowledge, the **DCT** set has never been used for studies of primate vocalizations. It is based on studies of human speech where DCT coefficients are useful for characterizing time-varying sounds, such as diphthongs. With only 7 dimensions, it is a minimal set characterizing the tonal contour of the call, its acoustic roughness and its duration. Adopting these three feature sets is intended to test which are the most efficient in a classification task, whether they are redundant or complementary, once feature correlation is accounted for by the classification procedure (see Section VI for details).

#### Automatic classification approaches and evaluation methodology

A multi-label classification task aims to assign observations described by a set of predictors to one of several predefined classes (call types or individual identities here). This task is treated here as a supervised learning task, in which a model can be trained on a set of examples with known classes, and then used to classify new ones. We chose a form of discriminant analysis (Permuted DFA or pDFA) as a baseline, as it is a widely used classification technique in the field of animal communication, including in individual identification tasks (Clink, et al., 2017; Mundry & Sommer, 2007; Favaro, et al., 2015, Keenan, et al., 2020; Lee, et al., 2006; Li, et al, 2017; Mathevon, et al., 2010; Oyakawa, Koda & Sugiura, 2007, among many others). We also implement three other supervised approaches, which can be described as ‘state-of-the-art’ in data science. SVM (Support Vector Machines) have been considered one of the best approaches for classification in the early 21^st^ century and are widely used in classification problems, including in ecology and ethology (e.g., Cheng, et al., 2012; Clink, Crofoot, & Marshall, 2019; Versteegh, et al., 2016; Dezecache, et al., 2021). xgboost is a classification technique derived from decision trees (Friedman, 2001) and is currently considered as one of the best methods (Chen, et al., 2012; Chen & Guestrin, 2016). Neural networks (NN) have been around for several decades, but their performances have improved dramatically in the last decade after the discovery of how to efficiently train deeper architectures (LeCun, Kavukcuoglu, & Farabet, 2010). Although they achieve by far the best performance today in computer vision and natural language processing, with models now involving up to hundreds of billions of parameters (e.g., Chowdhery, et al., 2022), these large networks require (very) large training datasets, which do not fit very well in the context of SUNG datasets, despite recent attempts (Leroux, et al., 2021). Instead, we will consider ‘shallow’ dense neural networks (two to four fully connected layers, including the output layer) well suited for small datasets and the size of our different sets of predictors, as they have proven effective in similar applications (e.g., Mielke & Zuberbühler, 2013; Robakis, Watsa & Erkenswick, 2018).

Each of our ‘state-of-the-art’ approaches involves the tuning of a number of hyper-parameters, which values can impact performance on a given dataset (Ramasubramanian & Moolayil, 2019). While the parameters are adjusted during the training phase – e.g., the values of the connections between neurons in a NN –, the hyper-parameters are not and have to be specified otherwise - e.g., the number of neurons in each layer of a NN. We have implemented a usual approach in machine learning, which is to consider not only a training set and a test set, but also a validation subset taken off the training set and used alongside the remaining part of the training set to find the optimal values of the hyper- parameters for the data at hand, in a process known as hyper-parameter tuning.

In order to evaluate the performance of our different classification techniques, we had to specifically consider the imbalance in the dataset. Indeed, not all metrics are appropriate to face this situation. In particular, the accuracy, which is easily interpreted, returns results biased towards the more represented classes. When only two classes are involved, it is common to consider instead the numbers of true/false positives and negatives, along with related metrics (sensitivity, specificity, F1, ROC AUC, PR AUC etc.). Such metrics can be adapted to multi-class classification, as in our case. We therefore considered three measures in addition to standard accuracy (which we kept for the sake of comparison with previous studies): multi-class log loss (a.k.a. cross-entropy), multi-class AUC and balanced accuracy:

- Multi-class **log loss** penalizes the divergence between actual and predicted probabilities - lower values are better. Log Loss differs from the two next metrics in that it considers probabilities rather than classification outputs.
- Multi-class **AUC** (Hand and Till, 2001) extends (two-class) AUC with two possible binarization strategies of the multiple-class problem: i) reducing it to several one-versus-all-others problems or ii) reducing it to several one-versus-one problems (results can be averaged in both cases). We adopted the second option while considering additionally the a-priori distribution of the classes to better address imbalance.
- The **balanced accuracy** (bac) is defined as the average of recall (a.k.a Sensitivity) obtained on each class. This addresses the issue of standard accuracy being biased by classes of different sizes. Balanced accuracy offers the advantage of being straightforward to understand, compared to log loss and AUC.

To assess whether our performance was significantly above chance, we introduced a random baseline consisting of 1,000 pairs of training and test sets with random shuffling of the predicted variable call type or emitter identity, depending on the task. For each pair, performance was measured and together led to a distribution under the null hypothesis, which is used to estimate *p*-values.

We further evaluated the importance of the different features used as predictors when classifying the calls, in order to detect whether some of them play a significantly larger role than others. We analyzed the features of the feature sets ‘Bioacoustic’ and ‘DCT’, but did not consider MFCC as there are too many features leading to very limited – and mostly meaningless – impacts for the individual features.

Different feature sets and classifiers may model differently the information present in the dataset. This suggests that they can be combined to achieve a better performance by cumulating their individual strengths while mitigating their individual weaknesses. So-called *ensembling* methods have been successfully developed for a large number of machine learning challenges. Bagging, boosting and stacking are three popular methods to combine different models (Bauer & Kohavi, 1999), depending on whether one builds models with different subsamples of the training dataset, one chains models to gradually reduce prediction errors (like in xgboost), or one integrates several parallel models applied to the same observations, which is the approach we implemented in this study. Different stacking approaches are available: simple ones like voting or averaging predictions, and more advanced ones which involve additional supervisor models - known as super learners - using the predictions of the initial models as inputs. xgboost may for instance be used as a super learner, and hyper-parameter tuning of the super learner may even be conducted. Stacking works best when several models with already good performances are combined.

Once an ensemble learner is defined, its performances can be evaluated exactly in the same way as non-ensemble ones, allowing one to estimate the gain of the ensembling strategy. We defined and implemented different stacked learners combining feature sets and classifiers (full description can be found in Section Methodology, Ensembling) and tested them along with the individual classifiers.

We mainly report results with balanced accuracy below, as it is more directly interpretable than log loss and AUC. More details about the approach can be found in Section VI (Methodology), as well as in Supporting Information. In particular, the full code to replicate the analysis is provided.

#### Task 1: Identification of call types

Our results (Figure 5 and Table 2) confirm that the five call types considered are discriminable to some extent, with a balanced accuracy reaching 0.794 with the best classifier.

**Figure 5.**
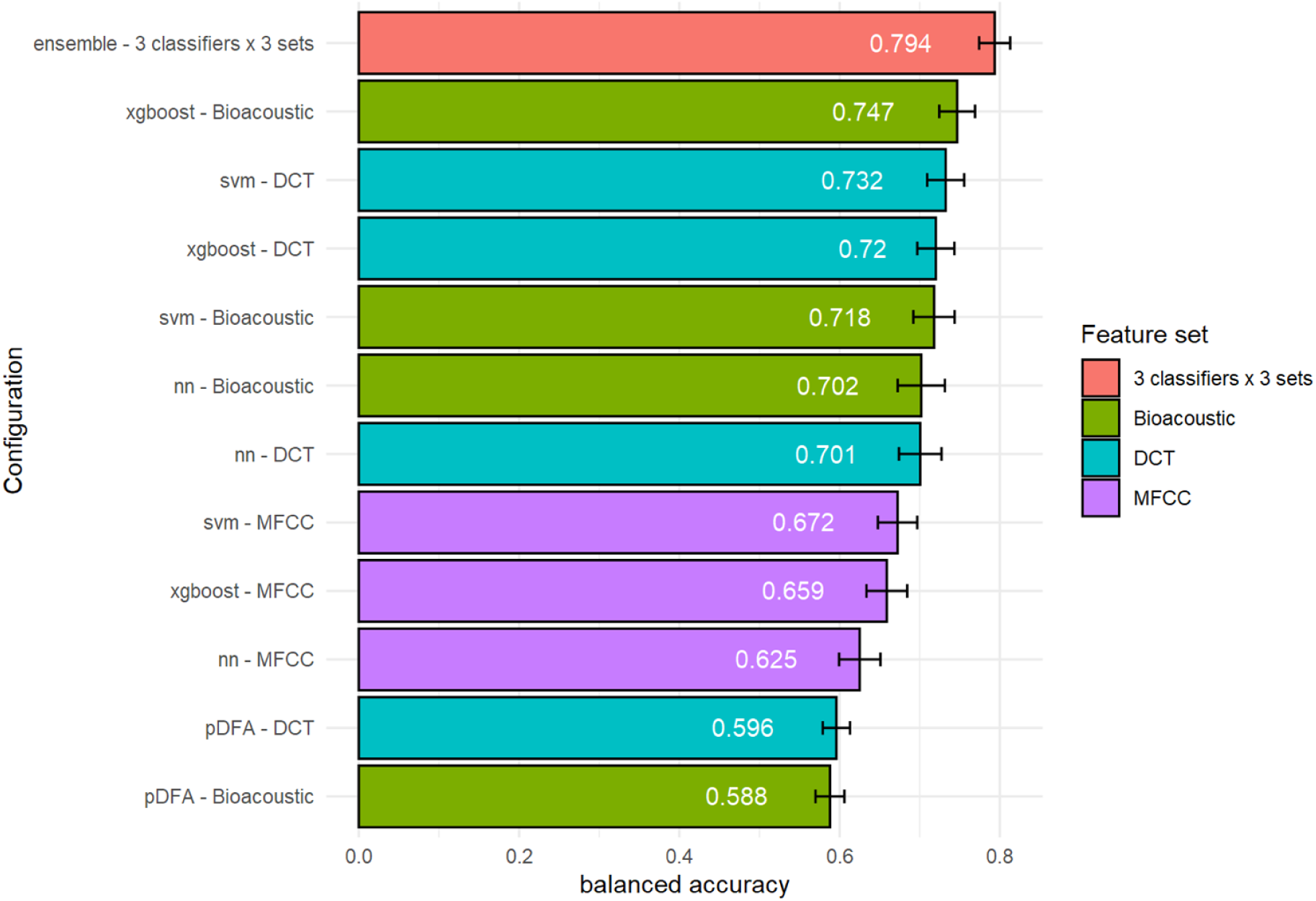
Performance in classifying bonobo call types as a function of classifier and acoustic set used. The red bar shows the performance achieved by an ensemble classifier combining the 9 primary classifiers. The other bars correspond to configurations associating each classifier with different sets of acoustic features (Bioacoustic, DCT, MFCC). The configurations are sorted by decreasing performance from top to bottom. Performance is reported in terms of balanced accuracy. Green, turquoise, and purple indicate the models trained on the Bioacoustic, DCT, and MFCC feature sets respectively.

**Table 2.** Metrics characterizing the classification performance of call types as a function of the classifier and acoustic set used. Four metrics are reported: log loss, AUC, balanced accuracy and accuracy. The best performance achieved by a primary configuration (upper part) and an ensemble configuration (lower part) is displayed in bold.

| Algorithm | Feature set | Config. # | Combined config. | log loss | AUC | accuracy | balanced acc. |
| --- | --- | --- | --- | --- | --- | --- | --- |
| pDFA | Bioacoustic |  |  | 0.819 | 0.894 | 0.651 | 0.588 |
| pDFA | DCT |  |  | 0.813 | 0.901 | 0.674 | 0.596 |
| svm | DCT | (1) |  | 0.658 | 0.934 | 0.745 | 0.732 |
| svm | Bioacoustic | (2) |  | 0.671 | 0.931 | 0.736 | 0.718 |
| svm | MFCC | (3) |  | 0.810 | 0.906 | 0.664 | 0.672 |
| nn | DCT | (4) |  | 0.681 | 0.932 | 0.724 | 0.701 |
| nn | Bioacoustic | (5) |  | 0.674 | 0.933 | 0.728 | 0.702 |
| nn | MFCC | (6) |  | 0.890 | 0.888 | 0.621 | 0.625 |
| xgboost | DCT | (7) |  | 0.651 | 0.936 | 0.732 | 0.720 |
| xgboost | Bioacoustic | (8) |  | **0.627** | **0.940** | **0.755** | **0.747** |
| xgboost | MFCC | (9) |  | 0.830 | 0.901 | 0.654 | 0.659 |
| ensemble |  |  | (1+2+3) | 0.640 | 0.939 | 0.759 | 0.744 |
| ensemble |  |  | (4+5+6) | 0.648 | 0.938 | 0.752 | 0.736 |
| ensemble |  |  | (7+8+9) | 0.792 | 0.912 | 0.675 | 0.683 |
| ensemble |  |  | (1+4+7) | 0.581 | 0.952 | 0.785 | 0.784 |
| ensemble |  |  | (2+5+8) | 0.604 | 0.950 | 0.773 | 0.767 |
| ensemble |  |  | (3+6+9) | 0.578 | 0.952 | 0.785 | 0.784 |
| ensemble |  |  | (1+2+3+4+5+6+7+8+9) | **0.542** | **0.957** | **0.794** | **0.794** |

The three classifiers SVM, NN, and xgboost outperform the pDFA approach, both with the bioacoustic set and the DCT set, and independently of the metric considered. The pDFA therefore partially misses some of the discriminative information available in the acoustic features. The balanced accuracy obtained with the pDFA is indeed only 0.596 with the bioacoustic set. This performance is comparable to that obtained by Keenan, et al. (2020) with the same method (57% of accuracy in a 5-category task with a slightly different call type labeling).

In order to compare the results of the discriminant analyses to the chance level, modified datasets were created by recombination (see Section VI Methodology) and a DFA applied to them. This 1,000-fold iterated procedure provided a robust estimate of the distribution of random accuracies. The empirical p-values obtained after this recombination procedure were equal to p=0.001.

Leaving aside stacked learners, it can also be seen that i) the results obtained with the MFCC set are worse than those obtained with the bioacoustic or DCT sets, ii) the best performances are obtained with the xgboost approach, while the worst are obtained with the NN approach. The best performing configuration is therefore the one that combines xgboost with the bioacoustic set. Although the MFCC set carries a richer description of the calls, using it does not bring any advantage, and even degrades the performance of the classifiers. The better performance achieved with the bioacoustic set is also consistent with the fact that the bioacoustic features are the cornerstone on which each call type is primarily defined by expert bioacousticians and primatologists. Finally, the fact that the performance reached with the DCT set is almost as good as with the bioacoustic set is very encouraging: it indicates that a small number of automatically extracted acoustic descriptors succeed in capturing most of the relevant information present in the signal.

When it comes to ensembling, all seven configurations improve the performance of the classifiers (or learners) they build upon. The best results are obtained with the stacking of all nine learners. The improvement is once again obvious, especially compared to the pDFA approach.

Furthermore, the observed difference between accuracy (acc) and balanced accuracy (bac) tends to be smaller for the stacked classifiers than for each algorithm separately, suggesting that the former handles class imbalance better.

Comparison to the random baseline showed that all results for the non-stacked learners - regardless of the feature set and the metric - are significantly above chance level with *p* < 0.001.

Focusing on the best performing approach – the stacking of the 9 different configurations – Figure 6 displays the average confusion matrix for 100 iterations. It confirms the quality of the classification, but also highlights a certain degree of confusion between several call types: between B and SB and between SB and PY, and to a lesser extent between B and SCB and between P and PY. Confusion thus occurs mainly between call types that are “adjacent” in terms of duration.

**Figure 6.**
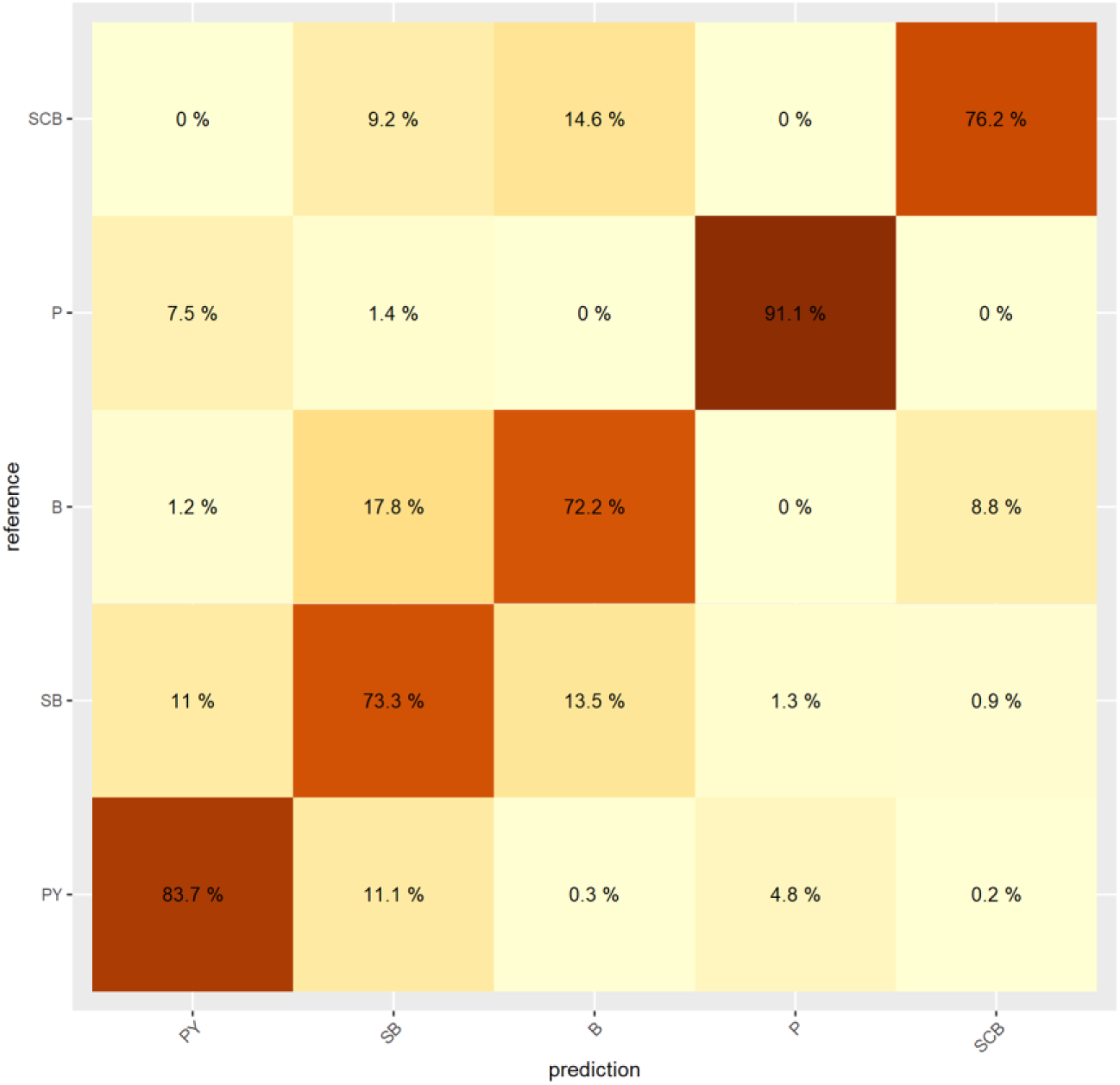
Confusion matrix reporting the classification rates of the call types in the best configuration (the ensemble classifier combining the 9 primary classifiers). Types are sorted from bottom to top by decreasing number of occurrences (PY: most frequent; SCB: least frequent). Percentages are according to the reference and sum to 1 along rows. The value of a cell color is proportional to its percentage (the darker, the larger).

Figure 7 shows the relative importance of the different acoustic descriptors as estimated with xgboost. Duration appears to be the most important feature, followed by f0.onset and f0.offset. For the DCT approach, dct2 – related to the curvature of the F0 trajectory – and duration are the two major predictors. The SVM and NN approaches give similar information. These results show that call types can be characterized by very few acoustic descriptors.

**Figure 7.**
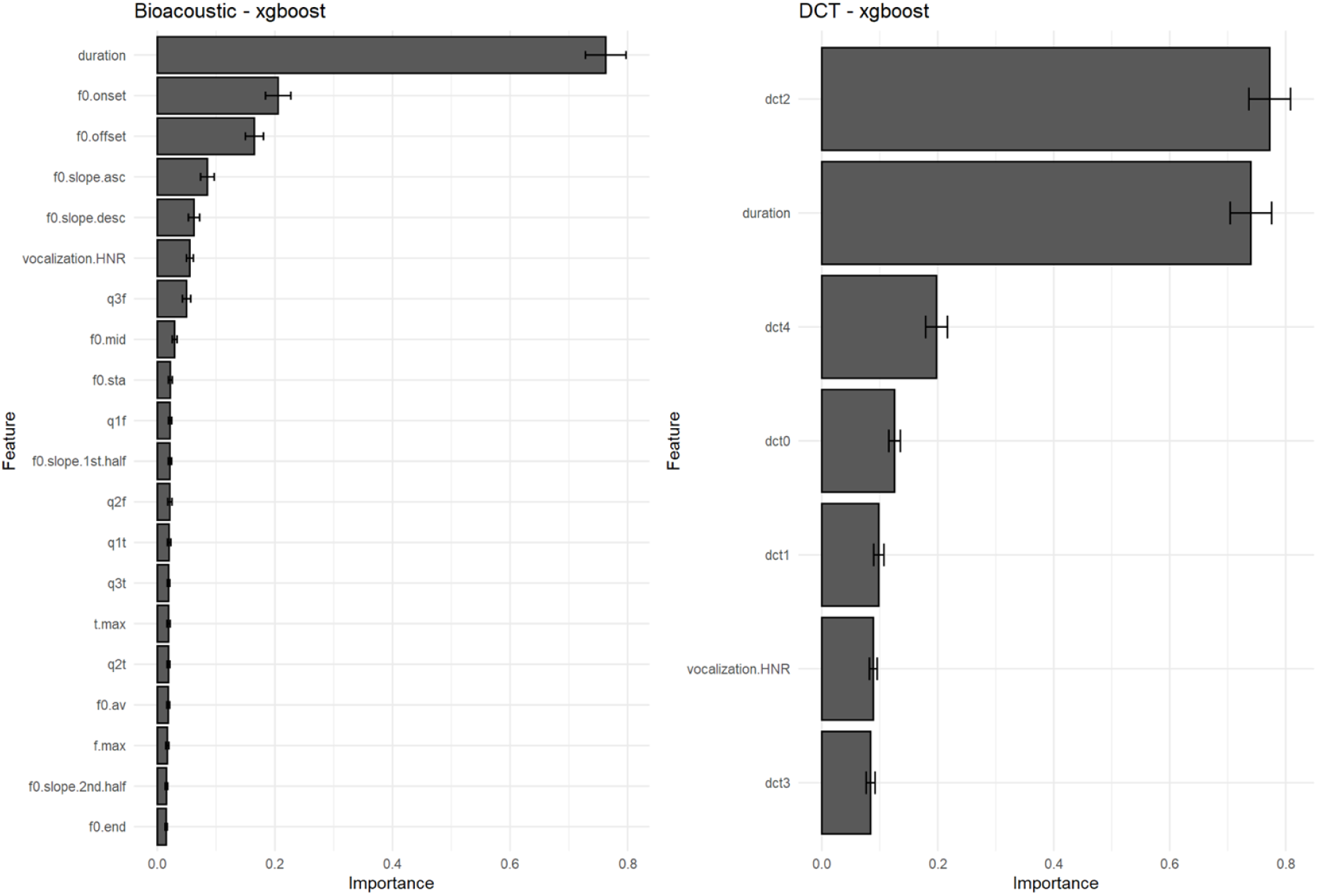
Importance of acoustic features when classifying call types with xgboost. **Left**. Features of the Bioacoustic set. **Right**. Features of the DCT set. The bar plots illustrate the relative influence of each acoustic feature on the classification performance.

#### Task 2: Identification of Individual signatures

Concerning the discriminant analyses applied to the individuals, the statistical significance of the accuracies was also higher than the chance level (empirical p=0.001). More generally, the best performance (bac = 0.507) in this 10-class problem is lower than for the 5-class call type classification. However, it is again much better than that given using pDFA (bac = 0.236). The difference in performance between the pDFA approach and the other approaches is even greater when it comes to identifying individuals than for call types (Figure 8 and Table 3). The three classifiers SVM, NN and xgboost again outperform pDFA with both bioacoustic and DCT sets, regardless of the metric considered.

**Figure 8.**
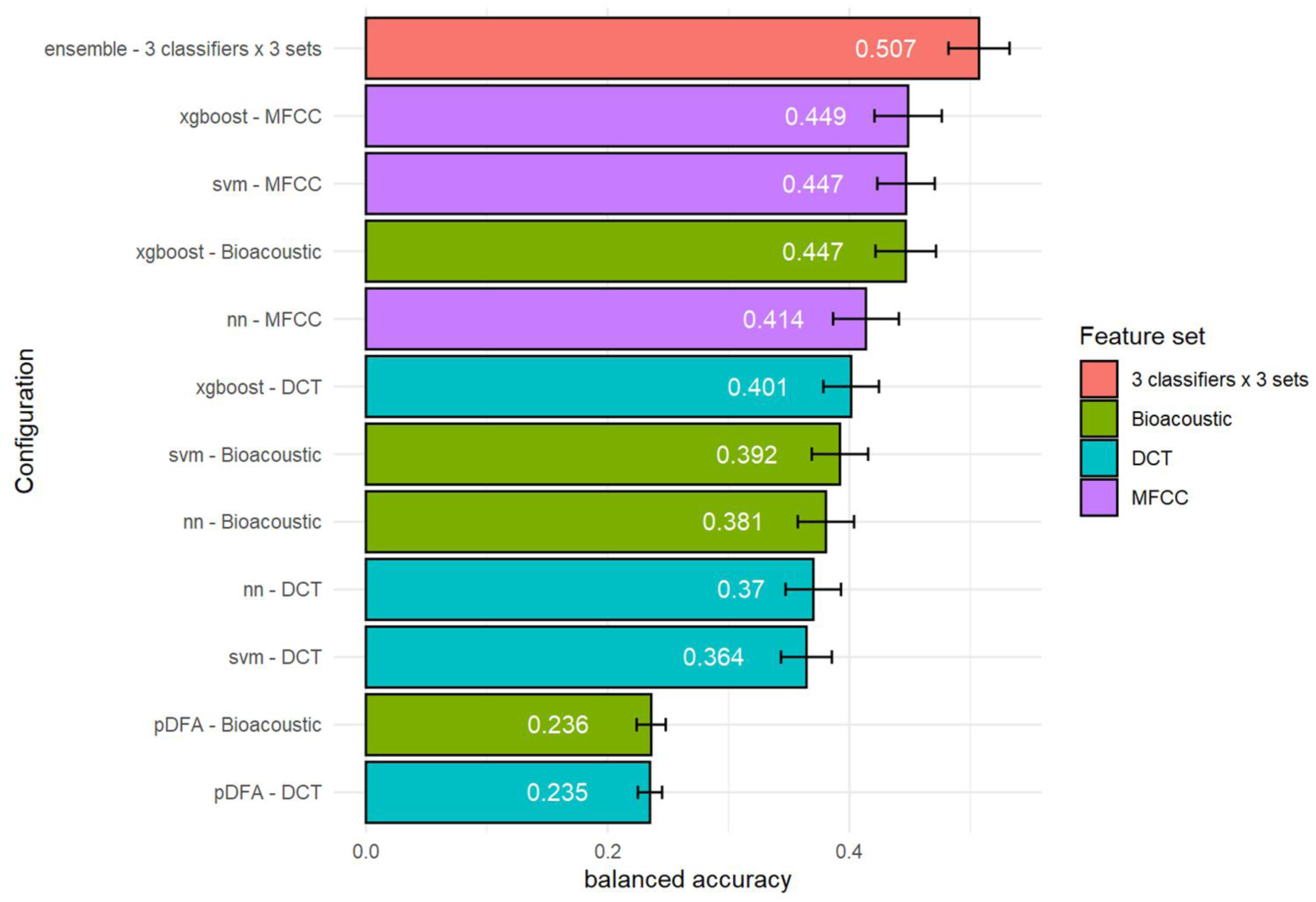
Performance in classifying bonobo individual signatures as a function of classifier and acoustic set used. The red bar shows the performance achieved by an ensemble classifier combining the 9 primary classifiers. The other bars correspond to configurations associating each classifier with different sets of acoustic features (Bioacoustic, DCT, MFCC). The configurations are sorted by decreasing performance from top to bottom. Performance is reported in terms of balanced accuracy. Green, turquoise, and purple indicate the models trained on the Bioacoustic, DCT, and MFCC feature sets respectively.

**Figure 9.**
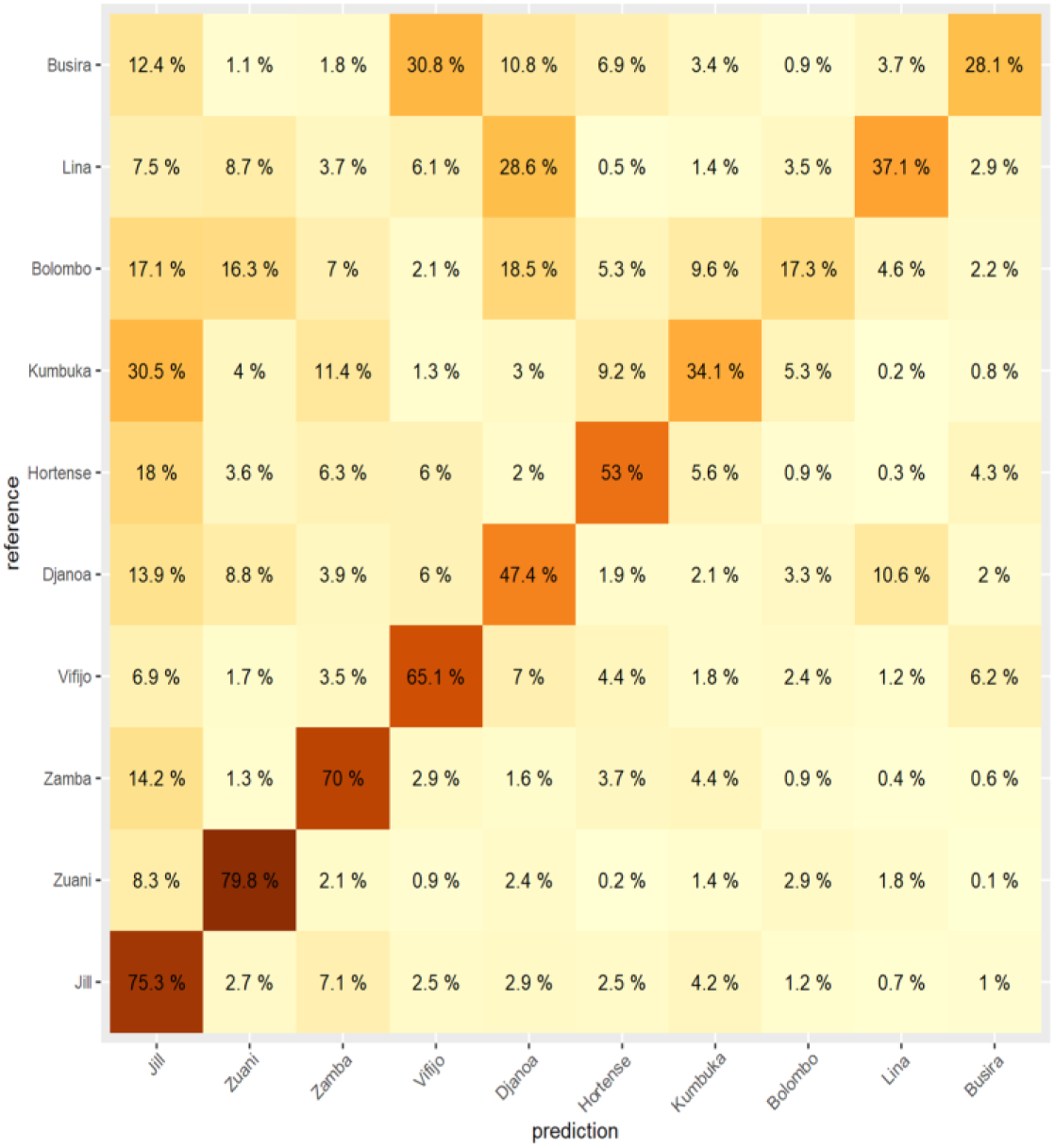
Confusion matrix reporting the classification rates of the individual signatures in the best configuration (the ensemble classifier combining the 9 primary classifiers). Individuals are sorted from bottom to top by decreasing number of calls (Jill: largest number; Busira: lowest number). Percentages are according to the reference and sum to 1 along rows. The value of a cell color is proportional to its percentage (the darker, the larger).

**Figure 10.**
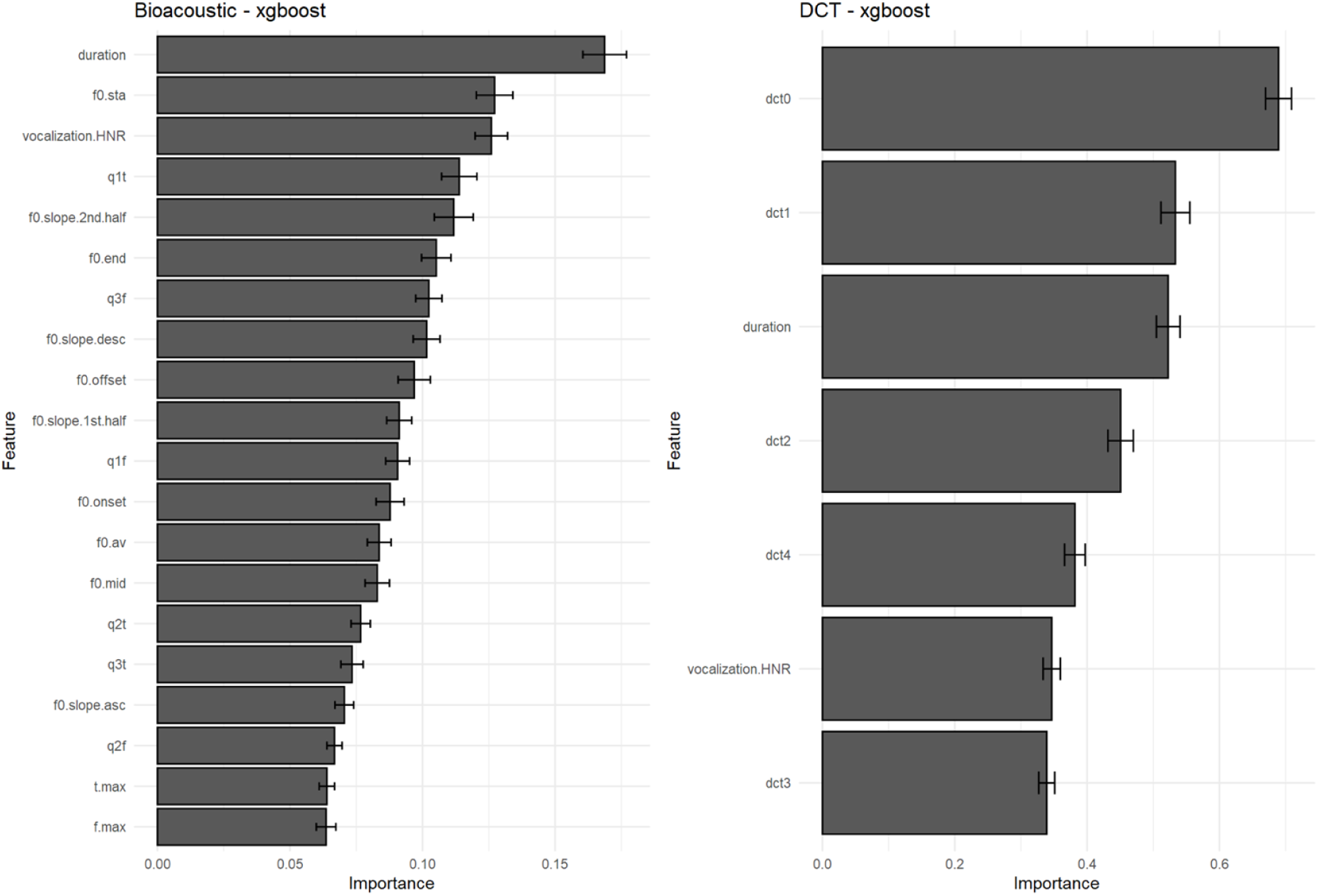
Importance of acoustic features when classifying individual signatures with xgboost. **Left**. Features of the Bioacoustic set. **Right**. Features of the DCT set. The bar plots illustrate the relative influence of each acoustic feature on the classification performance.

**Table 3.** Metrics characterizing the classification performance of individual signatures as a function of the classifier and acoustic set used. Four metrics are reported: log loss, AUC, balanced accuracy and accuracy. The best performance achieved by a primary configuration (upper part) and an ensemble configuration (lower part) is displayed in bold.

| Algorithm | Feature set | Config. # | Combined config. | log loss | AUC | accuracy | balanced acc. |
| --- | --- | --- | --- | --- | --- | --- | --- |
| pDFA | Bioacoustic |  |  | 1.745 | 0.731 | 0.410 | 0.236 |
| pDFA | DCT |  |  | 1.698 | 0.731 | 0.429 | 0.235 |
| svm | Bioacoustic | (1) |  | 1.490 | 0.840 | 0.516 | 0.392 |
| svm | DCT | (2) |  | 1.567 | 0.822 | 0.486 | 0.364 |
| svm | MFCC | (3) |  | 1.416 | 0.855 | 0.548 | 0.447 |
| nn | Bioacoustic | (4) |  | 1.487 | 0.839 | 0.508 | 0.381 |
| nn | DCT | (5) |  | 1.537 | 0.829 | 0.490 | 0.370 |
| nn | MFCC | (6) |  | 1.415 | 0.855 | 0.536 | 0.414 |
| xgboost | Bioacoustic | (7) |  | 1.415 | 0.850 | 0.544 | 0.447 |
| xgboost | DCT | (8) |  | 1.530 | 0.827 | 0.493 | 0.401 |
| xgboost | MFCC | (9) |  | **1.366** | **0.861** | **0.552** | **0.449** |
| ensemble |  |  | (1+2+3) | 1.240 | 0.887 | 0.594 | 0.495 |
| ensemble |  |  | (4+5+6) | 1.264 | 0.886 | 0.583 | 0.466 |
| ensemble |  |  | (7+8+9) | 1.254 | 0.885 | 0.589 | 0.484 |
| ensemble |  |  | (1+4+7) | 1.420 | 0.851 | 0.542 | 0.427 |
| ensemble |  |  | (2+5+8) | 1.493 | 0.835 | 0.500 | 0.389 |
| ensemble |  |  | (3+6+9) | 1.311 | 0.873 | 0.571 | 0.465 |
| ensemble |  |  | (1+2+3+4+5+6+7+8+9) | **1.210** | **0.894** | **0.605** | **0.507** |

Leaving stacked learners aside, the best performance is obtained with the MFCC set, then the bioacoustic set and finally the DCT. The best performing classifier approach is xgboost. The descriptors/classifier combination giving the best results is MFCC and xgboost. Contrary to what we found with the call types, the richness of the MFCC description enhances discrimination between the individual signatures. This result suggests that the bonobo vocal signature results from salient differences in the way each individual “positions” its calls (as illustrated in Figure 4 by the differences observed between the templates of individuals #19 and #20), complemented by subtle variations more easily captured by MFCC than by standard bioacoustic features.

When it comes to ensembling, six of the seven configurations improve the performance of the learners on which they are based while stacking the three algorithms NN, SVM and xgboost with the same set of bioacoustic features does not bring any improvement. This suggests a ceiling effect. The best results are obtained with the stacking of all 9 configurations.

As with call types, all classification results for the non-stacked learners - regardless of feature set and metric - are significantly above chance level with *p* < 0.001. However, the impact of the unbalanced dataset is striking. With the ensemble configuration leading to the best performance – the stacking of the nine different configurations – we obtain, on the one hand, quite good performances (up to 79.8% correct identification) for the four individuals contributing the most to the dataset (Jill, Zuani, Zamba, Vifijo). On the other hand, the performances are modest, though above chance, for the individuals that contribute less (e.g., Bolombo = 17.3% of correct identification; Busira = 28.1%). Class imbalance thus

has a significant impact on our results, despite the adoption of strategies to mitigate it. These results suggest that when a poor individual classification score is obtained, it is likely to be due to a faulty classifier and not to the absence of idiosyncratic features in an individual’s calls.

By examining the impact of each feature on classification performance, it can be observed that their importance is more diffuse across a wider set of features than was observed in the call type classification task, confirming the multidimensional and complex nature of the individual vocal signature.

### III. Addressing possible data leakage

#### Procedure

In the automatic classification approach reported in the previous section, each observation unit consists of a single call. A problem may arise when these calls are extracted from the same vocal sequence, which is frequently the case in animal datasets. This situation violates the independence of the observations (pseudoreplication), and can potentially undermine the validity of the classification performance. Specifically, how can we be sure that certain features characterizing the call sequence as a whole are not used by the classifier to identify the emitter? If this is the case, as suggested for instance by Clay and Zuberbühler (2011), single call classification performance could be overestimated.

To address this issue, we compared three different subsampling strategies to build training and test subsets. The first strategy (called *Default*) corresponds to the results reported in Section II. It simply consists in not exercising any control over how calls are assigned to one or the other subset, other than to ensure similar distributions of the occurrences of individuals in both sets. A second strategy (*Fair* strategy) consists in minimizing overlap (i.e. calls belonging to the same vocal sequence) by assigning as many sequences as possible to either the training set or the test set, so that the soundscapes of the sequences seen during training do not provide any information when classifying calls in the test phase. Full independence can be achieved in theory if enough data sequences are available to match the distributions of types and individuals between the training and test sets, but in practice the limited size of the dataset leads to residual leakage (see results below).

Finally, the third strategy (*Skewed* strategy) consists in maximizing the proportion of sequences shared by both sets (but still with disjoint call sets). By definition, the *Skewed* strategy is ill-designed as it maximizes data leakage, which automatically leads to an overestimation of classification performances. It is nevertheless instructive in providing ceiling performance against which the *Default* and *Fair* strategies can be compared.

## Results

To assess the influence of data leakage, we drew the training and test sets 100 times following each sampling strategy. The outputs of our approach are displayed in Figure 11. On the left side, the horizontal axis corresponds to a measure of the degree of overlap, defined as the number of call swaps required for all sequences to appear in a single set (ignoring the constraint of predefined size for both sets). The count value is thus equal to zero for an ideally fair split without overlapping sequences between the training and test sets. It can be seen that doing nothing (Default) is actually closer to maximizing overlapping (Skewed) than to minimizing it (Fair).

**Figure 11.**
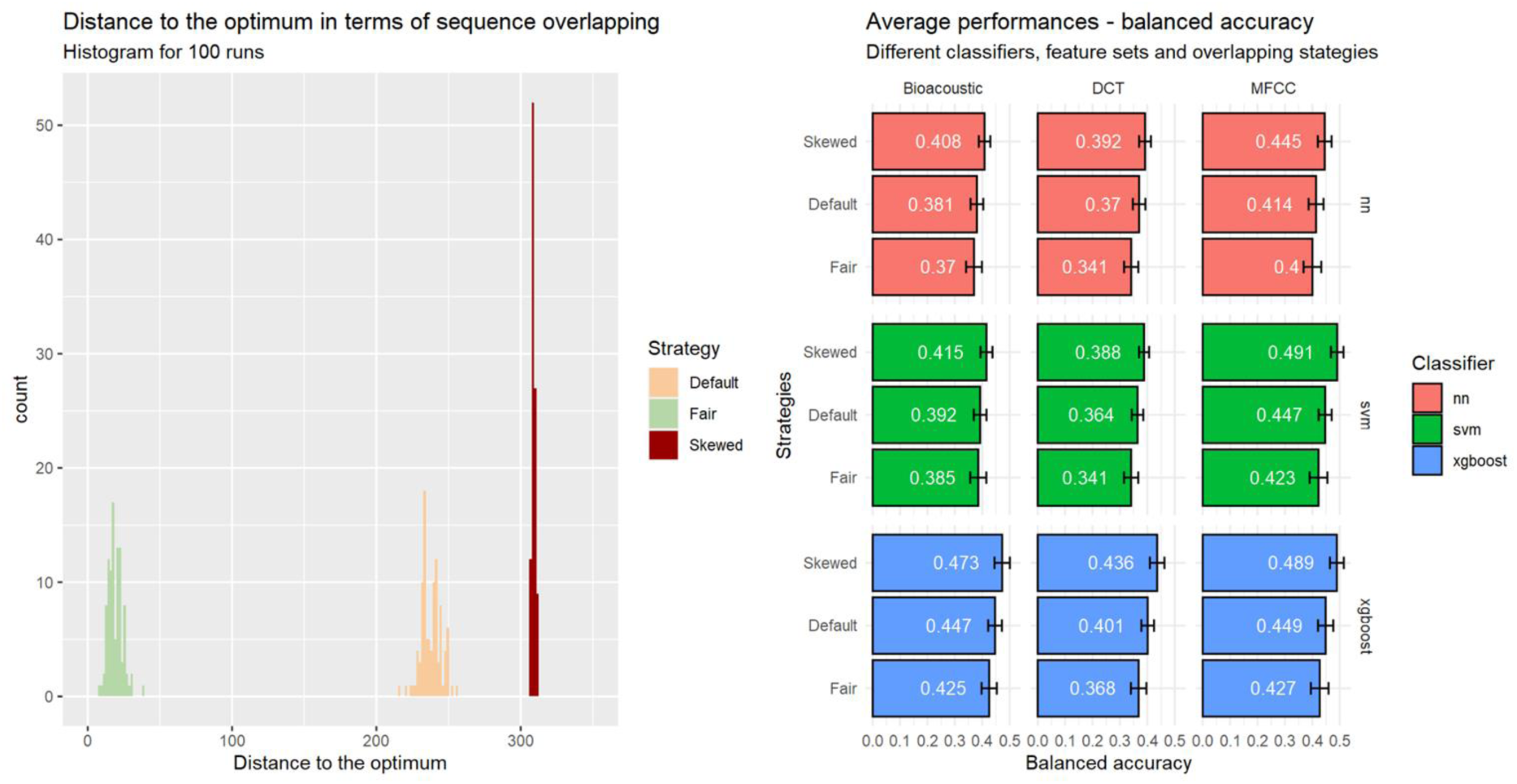
Influence of the sampling strategy on data leakage (all sequences considered). Three strategies are applied: *Default*, *Fair* and *Skewed*. **Left.** Distribution of the 100 runs for each strategy in terms of sequence overlap between training and test sets (0: no overlap). **Right.** Influence of strategy on performance (balanced accuracy) for each combination of classifiers and acoustic feature sets when classifying individual signatures.

We hypothesize that the performance would be the highest for the Skewed strategy and the lowest for the Fair one. In addition to the 100 runs reported in the previous section following the Default sampling, we computed 100 runs for both the Fair and Biased strategies. For the sake of simplicity, we left aside ensemble learners and focused on our 9 initial configurations.

The results can be found on the right side of Figure 11. Our hypotheses are confirmed, i.e., preventing overlapping of the sequences leads to reduced performances, when maximizing it leads to inflated ones. The former are more reliable as they correspond to minimization of the issue of non- independence between the observations where it matters most, i.e., between the training set and the testing set. One can observe, however, that the differences between the different strategies are small, which raises the question whether it is really necessary to control for the grouping of calls in sequences. Additionally, there are no differences in the general pattern of performances across classifiers and sets of predictors. Results are similar across our different metrics (see Supporting Information).

One explanation to the previous observations may lie in how calls are specifically organized in sequences in our dataset. Out of the 571 sequences in our dataset of 1,560 calls, 259 sequences consist of only one call, and 111 consist of 2 calls. This may explain why the differences between the different strategies are limited: by definition, a one-call long sequence cannot be shared by the training and test sets…

To further test this possibility, we built a subset consisting only of calls appearing in sequences of at least 3 elements – all 10 individuals were still present for a total of 1,079 calls in 201 sequences. We then followed the same approach as with our primary dataset, considering our three different strategies with 100 runs for each, after having estimated the best hyperparameters for our 9 different configurations of sets of predictors and classifiers. Figure 12 reports the results as in Figure 11. While the overall pattern is unchanged, one can notice on the left side of the figure that the Default strategy is very close to the Skewed one, meaning that the ‘simplest’ approach is close to the worst scenario in data leakage.

**Figure 12.**
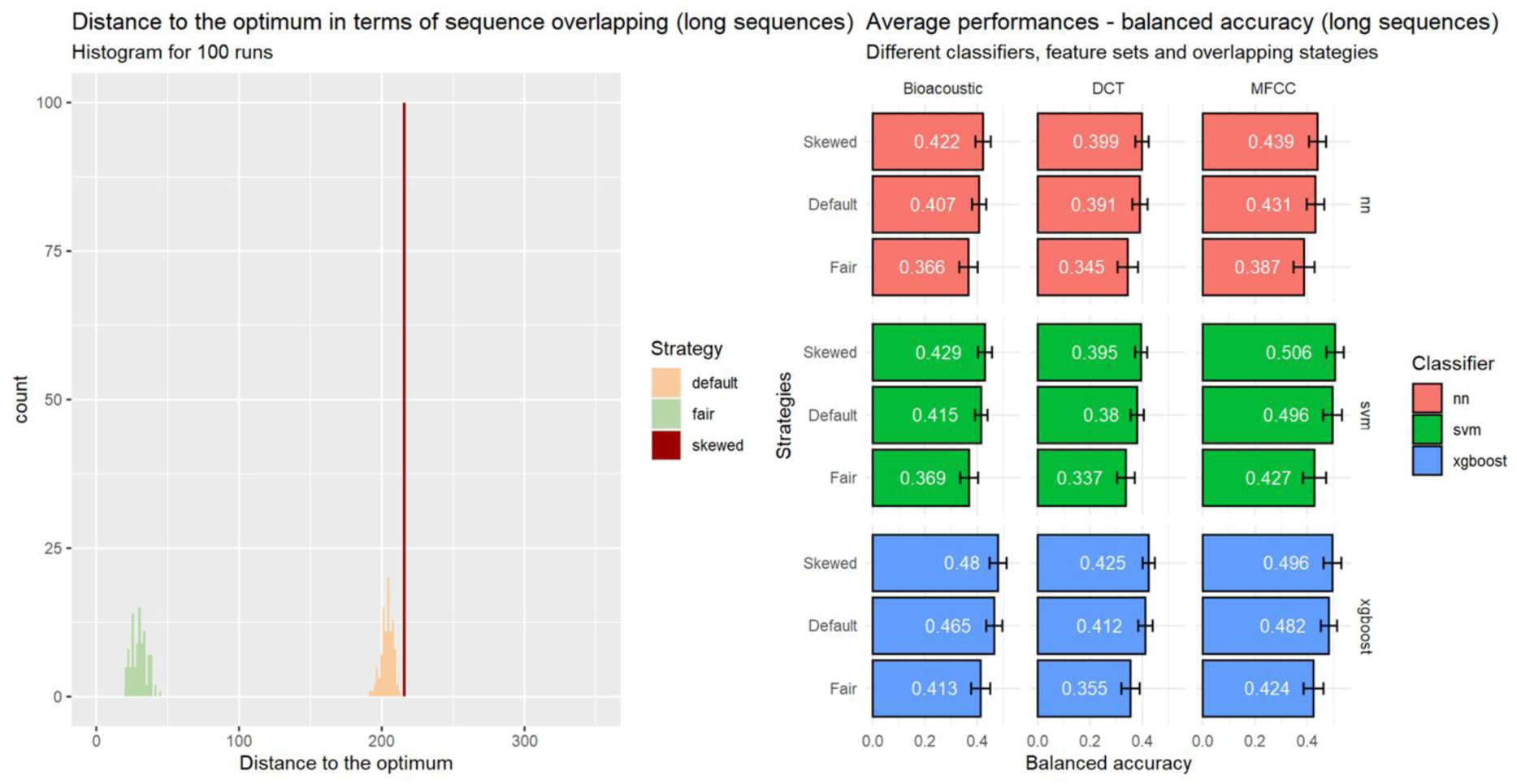
Influence of the sampling strategy on data leakage (sequences with at least three calls considered). Three strategies are applied: *Default*, *Fair* and *Skewed*. **Left.** Distribution of the 100 runs for each strategy in terms of sequence overlap between training and test sets (0: no overlap). **Right.** Influence of strategy on performance (balanced accuracy) for each combination of classifiers and acoustic feature sets when classifying individual signatures.

In terms of classification performance (Figure 12, right), one can observe larger differences between the Fair strategy and the others (e.g., a 12.5% gap in balanced accuracy between Fair and Default for xgboost with MFCC). It highlights that performance is clearly overestimated when the classifier can extract information that would be inaccessible in real life conditions.

The conclusion of our investigations is a cautionary tale: the occurrence of calls in sequences should be controlled to prevent the overestimation of classification performances, and this all the more as the average number of calls per sequence increases. The Skewed strategy should definitely be avoided since it dramatically overestimates the classification performances. On the contrary, one should try to implement a Fair strategy in order to get the least-biased performance estimation. If done manually, it can be tenuous, but optimization tools such as BALI provide an efficient way to alleviate the combinatorial burden. The Default strategy merely consists in the absence of any strategy and it is arguably a common practice when data leakage is – wrongly or rightly – not identified as a problem. One should mention that when data paucity is not an issue, practitioners often adopt a conservative approach that avoids most data leakage, for instance by systematically assigning recordings done on different dates to the training and test sets. Our caveat is thus mostly relevant in the context of SUNG datasets.

### IV. Comparing original vs. classifier-transformed representations

Confusion matrices provide an insightful overview of a classifier performance. Other approaches can additionally be considered to provide complementary assessments of the strengths and weaknesses of the classifier, along with a meaningful representation of data dispersion.

We considered UMAP (McInnes, et al., 2020) to visualize the performances of our different classifiers. We took advantage of the reduction of dimensionality offered by UMAP to contrast three different configurations for both individual signatures and call types: i) the raw highly-multidimensional description of the calls with the features of our three sets Bioacoustic, DCT and MFCC, ii) the predictions of pDFA with the Bioacoustic set, and iii) the predictions of the best ensemble. The resulting maps are found in the upper part of Figure 13 and Figure 14 for call types and individual signatures, respectively.

**Figure 13.**
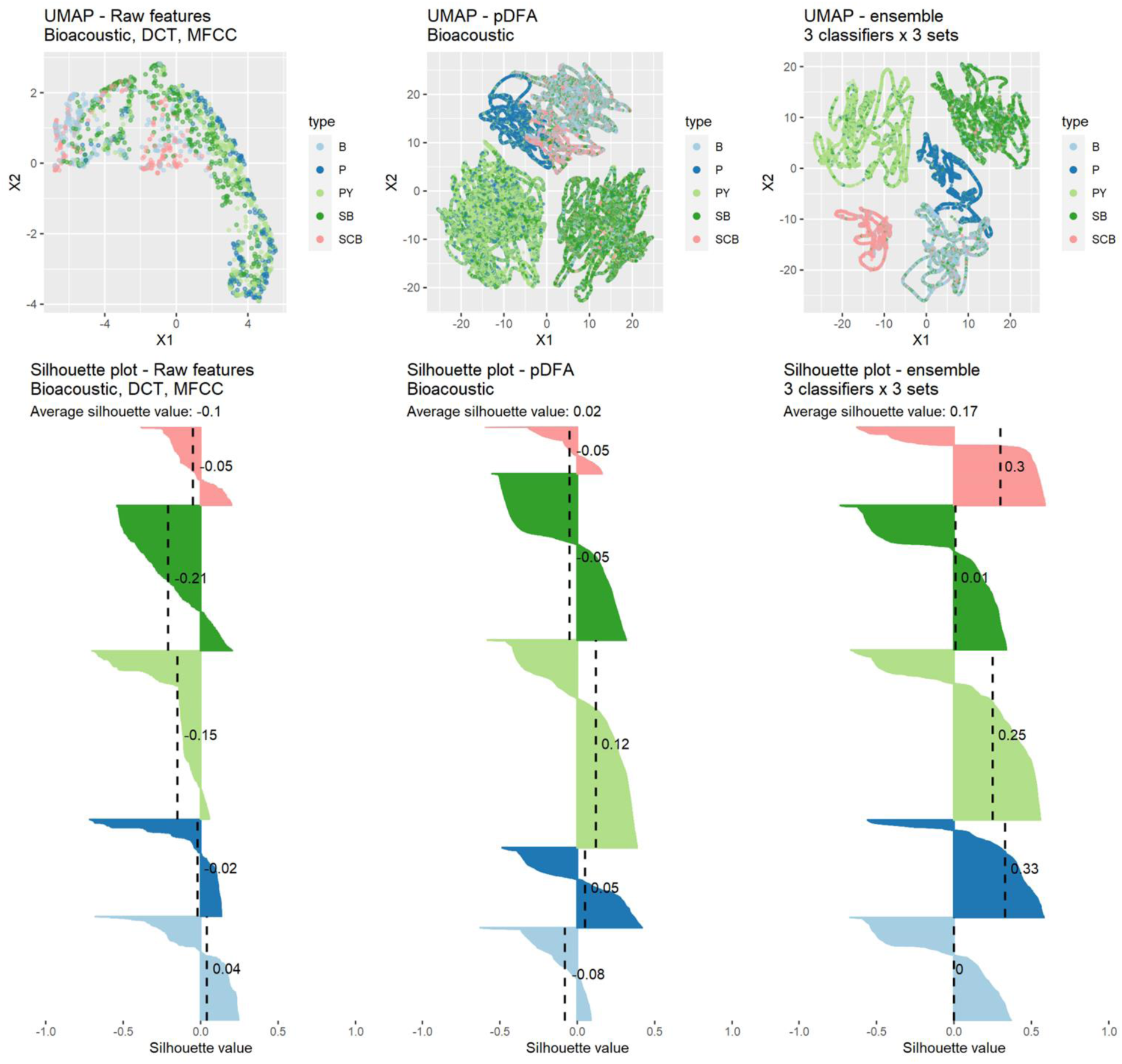
Bidimensional projections of bonobo calls into acoustic spaces with emphasis on call types (UMAPs; each dot = 1 call; different colors represent different call types). **Top, from left to right:** i) UMAP obtained with the raw acoustic features of the Bioacoustic, DCT, and MFCC sets (1,560 calls); ii) UMAP obtained with the outputs of a pDFA computed from the Bioacoustic set and the observations of our 100 test sets; iii) UMAP obtained with the outputs of the best ensemble classifier for the elements of our 100 test sets. **Bottom.** Silhouette profiles corresponding to the call type clustering. The overall average silhouette value and the average per call type are shown.

**Figure 14.**
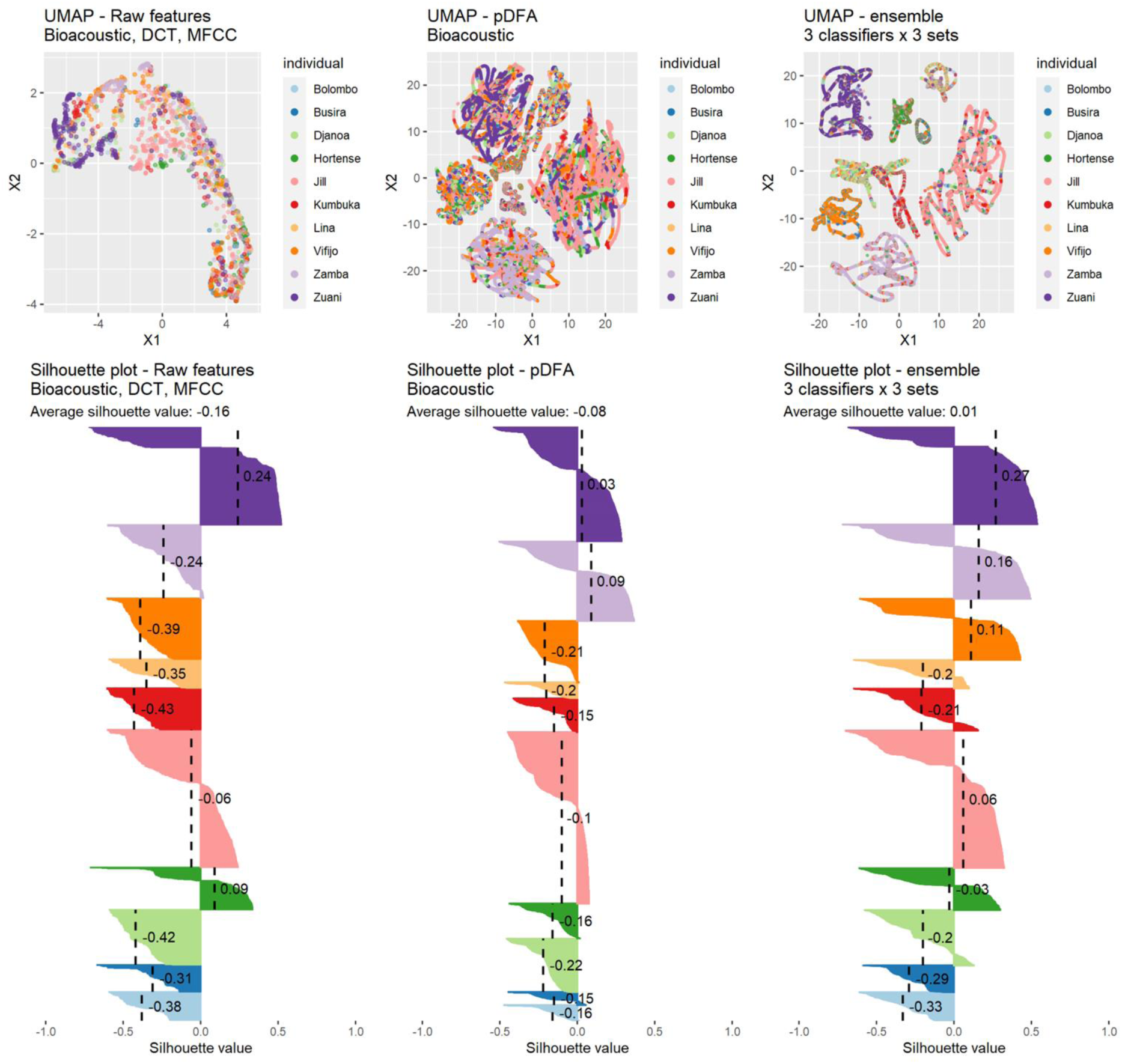
Bidimensional projections of bonobo calls into acoustic spaces with emphasis on individual signatures (UMAPs; each dot corresponds to one call, and one call can correspond to multiple dots - see more details in section VI; different colors represent different individuals). **Top, from left to right:** i) UMAP obtained with the raw acoustic features of the Bioacoustic, DCT, and MFCC sets (1,560 calls); ii) UMAP obtained with the outputs of a pDFA computed from the Bioacoustic set and the observations of our 100 test sets; iii) UMAP obtained with the outputs of the best ensemble classifier for the elements of our 100 test sets. **Bottom.** Silhouette profiles corresponding to the individual signature clustering. The overall average silhouette value and the average per individual are shown.

In order to estimate the quality of the partitioning provided by UMAP and thus quantify the degree of clustering of call types and individual signatures, we computed the values of the silhouette coefficients (Rousseeuw, 1987). For each data point *i*, *s*(*i*) is the difference between the average distance to data points in the same cluster and the average distance to data points in other neighboring clusters. The values of *s*(*i*) range from -1 to 1. A large positive coefficient indicates that a data point is close to elements in the same cluster, a small positive coefficient denotes a data point close to the decision boundary, while a negative coefficient indicates that it is closer to elements in another cluster. We computed i) the overall average silhouette score over the whole dataset (overall average silhouette width) ii) the average silhouette score per class (average silhouette width). The outputs for the three configurations are found in the lower part of Figure 13 and Figure 14, respectively for call types and individual signatures.

Visual clustering of either call types or individual signatures is improved when one shifts from raw features to pDFA to the representation derived from the best ensemble classifier. This is confirmed by the increasing values of overall average silhouette values from one configuration to the other. Additionally, in the case of the best ensemble, silhouette shapes and values for the different groups are consistent with the confusion matrices in Figure 6 and Figure 9. For the five call types, the Spearman’s rank-order correlation coefficient *ρ* between the percentage of correct classification (Figure 6) and the average silhouette values per group (Figure 13) is 0.9. For the 10 individuals, the coefficient of correlation is even higher: *ρ*=0.96. This clear correlation highlights that UMAP and silhouettes are consistent with our other assessment of the performances of our classifiers and furthermore that it is possible to get a bidimensional representation that accounts for most of the discriminability revealed by the classifier. Nevertheless, it should also be noticed that the silhouette scores for all the classification methods were below 0.2, which is rather low. It underlines that, while a variable proportion of the calls is distinguishable without any ambiguity, others belong to a gray zone where the classifier is indecisive in assigning a call type or an individual identity.

### V. Main achievements and limitations of the approach

While improving our understanding of how information is encoded in the bonobo vocal repertoire, this paper has primarily a methodological objective. Our goal is to assess the suitability of different approaches to vocalization processing (acoustic analysis, automatic classification and representation) to describe information encoded in vocalizations from a SUNG dataset representative of what is commonly obtained in bioacoustic studies (Figure 1). Our approach consists of an acoustic characterization in several feature spaces followed by the implementation of several classification algorithms and their combination to explore the structure and variation of bonobo calls (Figure 5) and the robustness of the individual signature they encode (Figure 8). We assess the importance of the notion of data leakage in the evaluation of classification performance, and the need to take it into account (Figures 11 & 12). Finally, we show how classifiers can generate parsimonious data descriptions that help to understand the clustering and structure of the acoustic space through a nonlinear UMAP representation (Figures 13 & 14).

#### Main findings

##### Features describing acoustic signals

We compared a set of 20 acoustic parameters traditionally used to describe primate vocalizations (Bioacoustic set) with a reduced set of seven semi-automatically computed parameters (DCT set) and a more comprehensive spectro-temporal parameterization comprising 192 parameters (MFCC set). While traditional acoustic parameters appear adequate for characterizing call types, it appears that classifiers benefit from a finer-grained acoustic description (such as that provided by the MFCC) when characterizing individual signatures. This is probably because the MFCC parameterization encodes subtle differences related to the emitter’s anatomy and physiology beyond the overall basic characteristics of the calls. These parameters are probably sufficiently idiosyncratic to allow an ape to correctly identify the emitter in a real life situation. An algorithm-based automatic classification is of course only partially related to the cognitive task an animal performs when it has to decode the information carried by a conspecific’s vocalization (Fedurek, Zuberbühler & Dahl, 2016; Linhart, et al., 2022), but it can be assumed that individual conspecific individual vocal identification is quite accurate in bonobos. This has been shown in an experimental study using playbacks with a single call type (Keenan, et al., 2016).

The fact that the performance reached with the DCT set on call type identification is only slightly worse than the best performance suggests that this representation adequately captures their essential aspects. This is very encouraging as it can be expected that automatic feature extraction can be rapidly developed for bonobo (and probably other mammals) calls, also taking advantage of recent improvements in acoustic pre-processing (Knight, et al., 2020; Stowell, et al., 2019; Stowell, 2022). Adopting multiple feature sets may also be good practice for extracting non-redundant information, particularly in small datasets. The complementarity between predefined acoustic features (Bioacoustic or DCT set) and agnostic spectro-temporal representation (MFCC set) echoes the conclusions of Sainburg et al. (2020) in the context of unsupervised visualization of a vocal repertoire.

##### Automatic classification

We implemented three classifiers (svm, xgboost, and NN) and compared their performance to that of discriminant analysis (pDFA), a method classically used to analyze the information carried by animal vocalizations, in particular vocal signatures. Our results show that all three models significantly outperform pDFA for the identification of both call types and individual signatures. Because it may not be sensitive enough, pDFA may miss the identification of acoustic categories: it is therefore not the most adequate means to assess the categorical nature of a complex vocal repertoire or the accuracy of vocal signatures. Furthermore, the performance achieved when evaluating individual signatures (illustrated by the confusion matrix in Figure 8) shows a striking contrast between individuals for which the dataset is very limited (around ∼75 calls for each of the three least represented individuals) and the most represented individuals (∼200 calls or more for three individuals). This difference reminds us that we have processed a very small dataset (Table 1) by the standards usually expected for machine learning. Moderately increasing the dataset size by only a few hundred observations per individual would certainly bring a significant improvement in the performance of the classifiers.

From a data scientist’s point of view, it is also interesting to note that ensembling methods improve performance. This means firstly that the three feature sets encode complementary information to some extent. Furthermore, the three classifiers exploit them in different ways. Obviously, bioacousticians are not necessarily obsessed with achieving the best classification performance. In fact, a trade-off between performance and computational time should in principle be sought. A simple classifier such as an svm can thus be sufficiently informative with the advantage of being simpler to implement and much less computer-intensive than xgboost for instance (especially when tuning the different hyper-parameters and in the repeated procedure necessary for a well-controlled cross-validation framework). High- accuracy classifiers could nevertheless be very useful when high accuracy is expected, for example when examining a new vocal repertoire, or for passive acoustic monitoring of wild animals (Gibb, et al., 2019; Kvsn, et al., 2020, Mcloughlin, Stewart, McElligott, 2019, for recent overviews).

**Performance assessment** must be done with great care. Dealing with a SUNG dataset means taking into account data sparsity, class imbalance, and potential data leakage. Their detrimental effects can be mitigated to some extent, by adopting the right metrics and models, and by carefully controlling for leakage induced by confound factors between the training and test sets. The high level of robustness observed in speech or speaker recognition with very large corpora (based on thousands of speakers and various recording conditions, to mention only two aspects) is unreachable with relatively small datasets. Even if one carefully assigns calls recorded at different times to the training and test sets, there is inherently some residual data leakage when assessing the individual signature, as the soundscape, recording conditions and equipment often vary between recordings. These caveats make it difficult to compare approaches and papers. In this regard, multicentric challenges offering a unified evaluation framework based on a shared dataset should be encouraged (Stowell, et al., 2019).

**Visualization.** Whether using illustrative raw spectrograms – as in Bermejo & Omedes (1999) and de Waal (1988) – or aggregated templates (as in this paper in Figures 3 and 4), graphical representations of vocalizations are essential to capture the structure and characteristics of any vocal communication system. The projection of a dataset into a feature space provides additional information about its variability and clustering. Recent nonlinear reductions and projections such as UMAPs are particularly convenient and powerful, as convincingly stated by Sainburg et al. (2020) for large datasets. However, these authors acknowledged that in the context of small datasets, UMAP representations may fail to clarify the structure of the dataset. Here, we overcome this limitation by showing that classifiers provide an elegant means of transforming the dataset from its acoustic description into an informative, parsimonious and discriminative latent feature space. This intermediate representation can in turn be used as a preprocessing step to better account for the clustering and discriminant potential of the data (see also Thomas, et al., 2022).

#### Recommendations

Our results lead to the identification of several practical approaches that are generalizable to any other animal communication system and can improve the standard bioacoustics workflow in terms of information gain and reproducibility, especially with SUNG datasets. These approaches may probably seem standard to some machine learning experts (Stowell, 2022), but the current literature shows that they are not yet adopted in a systematic way. However, this is not a magic recipe, and some thought should be given to whether these recommendations are relevant to the context of interest.

We recommend:

1. comparing several acoustic parameterizations as they may be complementary;
2. adopting SVM (Support Vector Machines) rather than discriminant functional analysis as the baseline classification approach;
3. explicitly evaluating data leakage and possibly implementing a mitigation strategy;
4. visualizing the dataset with a Uniform Manifold Approximation and Projection based on the classifier predictions rather than the raw acoustic features.

#### Future work

The approaches implemented in this study offer a reasonable balance between performance and complexity. Several recent approaches offer interesting alternatives or potential improvements. The direct use of spectrograms, either as an image or as a parameter matrix, has already been applied to mice (Premoli, et al., 2021), Atlantic spotted dolphins (Kohlsdorf, Herzing, & Starner, 2020), domestic cats (Pandeya, Kim, & Lee, 2018), common marmosets (Oikarinen, et al., 2019), etc.. However, their performance on complex and graded repertoires remain to be evaluated, and adaptation of spectrogram parameters to each species may be necessary (Knight, et al., 2020). Leroux et al. (2021) have recently fine-tuned a deep learning model trained on human voice to detect chimpanzee vocal signatures. This procedure is promising because conceptually it should work, but the results need to be confirmed on a larger task with stringent control of data leakage. Other neural network architectures have also been successfully implemented on large datasets (e.g. Ivanenko, et al. (2020) on mouse ultrasonic vocalizations). Progress has also been achieved in segmenting long audio bouts into calls (Oikarinen, et al., 2019; Steinfath, et al., 2021; Tachibana, et al., 2020), but the proposed solutions currently struggle with the adverse conditions typical of SUNG datasets (overlapping calls, non stationary noise, etc.). All these approaches are nevertheless expected to improve classification accuracy on large-enough datasets but the main performance limitation comes from the dataset itself. As aforementioned, its low audio quality may partially be compensated by an adequate acoustic preprocessing but complementary approaches lie in its size artificial extension by data augmentation techniques (Kahl, et al., 2021) or by developing multi-species models or transfer learning approaches (Provost, Yang, and Carstens, 2022). These techniques are not magic bullets, though, and their efficient implementation will require additional studies (Premoli, et al., 2021).

Finally, the automatic analysis approaches discussed in this paper provide efficient tools to help bioacousticians grasping a vocal repertoire. The proposed workflow addresses an intermediate, but essential, phase on the way to a global understanding of an animal communication system, and it thus takes part in a collective effort aiming at adapting recent machine learning approaches to bioacoustics.

### VI. Detailed methodology

#### Ethics statement

All research conducted for this article was observational. All data collection protocols were performed in accordance with the relevant guidelines and regulations and were approved by the Institutional Animal Ethical Committee of the University of Saint-Etienne, under the authorization no. 42-218-0901- 38 SV 09.

#### Extraction of acoustic features

- **Bioacoustics**: this set of acoustic parameters is inspired by the procedure described in Keenan, et al. (2020). It consists of parameters computed with Praat that summarize the F0 shape and the energy distribution for each call. The call harmonicity (or Harmonics-to-Noise Ratio) has also been included. By splitting the signal energy into its harmonic vs noise components, harmonicity takes the call acoustic roughness into account. This procedure results in 20 features. *Manual operations required: call segmentation and F0-Peak positioning*.
- **MFCC:** this set computed with Matlab is developed from a Mel-frequency cepstral analysis computed through 32 triangular shaped filters distributed over the 500-12000 Hz band. The coefficients are computed for successive frames (∼23 ms duration and 50% overlap) and a Hamming window is applied^1^. First and second order derivatives (so-called delta and delta-delta coefficients) are also computed, resulting in 3 x 32 = 96 dimensions. The final MFCC set consists of the average and standard deviation computed over the call. This procedure results in 192 features. *Manual operation required: call segmentation*.
- **DCTHarm**: this set is computed with Praat and is mainly based on the F0 contour, on which a Discrete Cosine Transform (DCT) is applied. DCT is a method used in phonetics to parametrize the contours of the fundamental frequency (F0) and formants (Zahorian & Jagharghi, 1993; Watson & Harrington, 1999; Teutenberg, Watson, & Riddle, 2008). This parametrization decomposes the signal into a set of ½ cycle cosine waves, the resulting amplitudes of these cosine waves being the DCT coefficients. The first DCT coefficient is a value proportional to the mean of the original F0 trajectory calculated between the onset and offset of the vocalization; the second coefficient is directly proportional to the linear slope of the F0; the third is related to its trajectory’s curvature. Each successive coefficient corresponds to a function with a ½ wavelength higher than its predecessor. As the number of coefficients used increases, the contour represented by the DCT coefficients approaches the original F0 contour. In the following analysis, the F0 contours have been approximated by the first five DCT coefficients. The call duration and harmonicity were also included. The formula used in a Praat script to compute the DCT coefficients was from Watson & Harrington (1999). This procedure results in 7 features (5 DCT coefficients + Duration + Harmonicity). *Manual operations required: call segmentation and F0 contour checking*.

All analyses detailed in the following methodological presentation were implemented with the R software (R Core Team, 2022). A detailed list of the packages is provided at the end of the section.

#### Data selection and preprocessing

We selected a subset of individuals and call types from our initial dataset, specifically the five types P (peep), PY -peep-yelp), SB (soft bark), B (bark), and SCB (scream bark), and individuals with more than 70 calls in these five types, namely Bolombo, Busira, Djanoa, Hortense, Jill, Kumbuka, Lina, Vifijo, Zamba, and Zuani. This selection resulted in a data set of 1,560 observations (Table 1).

One missing value was detected for f0.mid. A random forest algorithm trained on the observed values of the variables related to f0 (q1f, q2f, q3f, f.max, q1t, q2t, q3t, t.max, f0.sta, f0.mid, f0.end, f0.av, f0.max, tf0.max, f0.slope.asc, f0.slope.desc, f0.slope.1st.half, f0.slope.2nd.half, f0.onset, f0.offset) offered a plausible reconstruction.

Prior to the application of the classifiers, the different sets of predictors were standardized by centering them at the mean values and rescaling them using the standard deviation (z-scoring).

#### Imbalance in the data

Our dataset is characterized by an imbalance in the way our observations - the calls - are distributed across individuals and call types (not to mention other dimensions such as the context of emission). Table 1 summarizes the situation.

While the most represented individual - Jill - has 362 calls, the least represented individuals - Busira and Bolombo - have about 20% of this amount. The call types are more evenly distributed, with the number of occurrences ranging from 206 to 443. It is important to note that some individuals lack certain call types, e.g., Kumbuka and SCB, or have only a few occurrences, e.g., six B for Bolombo or five SB for Busira.

Such imbalance is a common feature of SUNG datasets, and should be taken into account when considering classification approaches. Although pDFA is inherently designed to deal with this type of imbalance, we considered several options for the other approaches. Results are reported with class weights inversely proportional to the number of observations in each class, which yielded better results than undersampling, oversampling or SMOTE (synthetic minority oversampling technique).

It should be noted that we only paid attention to the imbalance for the target domain of classification, i.e., when classifying individuals, we only considered the different numbers of calls per individual and not for the call types, and vice versa.

#### pDFA

In the first step of the permuted form of discriminant analysis, a training sample was used to compute a set of linear discriminant functions. In order to test for call type distinctiveness, the training sample was randomly selected from each call type without controlling for the individual. Similarly, in order to test the individual vocal signature, the training sample was randomly selected from each individual’s vocalizations. The number of occurrences selected was the same for each call type and each individual. This number was equal to two thirds of the smallest number of vocalizations analyzed per call type (i.e., 206×2/3) and per individual (i.e., 71×2/3). These discriminant analysis models computed from training samples were used to classify all remaining occurrences. Thus, for each individual and for each type of vocalization, at least one third of the occurrences were included in the test sample. We performed 100 iterations of these discriminant analyses with randomly selected training and test samples. The performance metrics and the percentage of correct classification were obtained by averaging the results obtained for each of the 100 test samples.

In order to statistically compare the performance achieved by the discriminant analysis to the chance level, additional discriminant analyses were also performed on modified datasets created by recombination, following the permuted DFA paradigm. Recombination involves assigning a class label (call type or individual identity, depending on the task under consideration) to an observation by a random permutation on the dataset. The training and test sets were then drawn according to the same rules as for the original dataset, and a DFA was applied, leading to an accuracy characteristic of a random classification. This procedure was iterated 1,000 times, resulting in a robust estimate of the distribution of random accuracies. The empirical p-value of the performance achieved with the original discriminant analysis was then obtained by dividing the number of randomized datasets that revealed a percentage of calls or individuals correctly classified at least as large as that obtained with the original dataset by the total number of datasets tested (Mundry & Sommer, 2007; Keenan, et al., 2020).

In contrast to the state-of-the-art approaches mentioned below, discriminant analysis does not automatically handle multicollinearity between predictors, which must be explicitly addressed. We chose to identify and reduce collinearity by computing the variance inflation factor (VIF). As suggested by Zuur, Ieno, & Elphick (2010), we used a stepwise strategy^2^. The VIF values for all predictors were computed, and if they were larger than a preselected threshold (here, 5), we sequentially dropped the predictors with the largest VIF, recalculated the VIFs and repeated this process until all VIFs were below the preselected threshold. A tick in Table A1 indicates the predictors included in pDFA analyses. Thus, since the assumptions of the discriminant analysis are stricter than those for state-of-the-art approaches, the MFCC predictor set was not used with the pDFAs due to its high dimensionality and collinearity which do not meet these assumptions (the size of the smallest class must be larger than the number of predictors).

#### Training/testing procedure

When working with a large dataset and considering a supervised ML technique, it is common to split it once between a training set and a test set. However, as the size of the dataset decreases, however, this binary sampling procedure gives rise to an increasing risk of sampling error. In other words, the results may depend significantly on the specific observations falling randomly into the two sets, as some observations are inherently more difficult to classify than others. It is therefore not prudent to assess performance with a single split^3^. To account for this risk, we averaged performances over 100 repetitions, i.e., over 100 pairs of training and test sets. We rely on 25-thread parallelization to reduce the time needed to obtain the results. The performance distributions clearly illustrate the effect and range of sampling error, and the overall symmetry of these distributions validates the use of mean values as an estimator of central tendency. We computed mean values not only for our different metrics (see below), but also for the confusion matrices and to assess feature importance.

#### Hyper-parameters for SVM, NN and XGBboost

We considered the following hyper-parameters for our different ‘state-of-the-art’ classifiers (with corresponding possible value ranges):

For svm: i) the nature of the kernel (linear, polynomial or radial), ii) for the polynomial kernel, the number of degrees (between 1 and 4), iii) the cost parameter C, which is a regularization parameter that controls the bias–variance trade-off (values between 2^-7 and 2^7, with a power-2 transformation), and iv) for the radial kernel, the parameter gamma or sigma, which determines how the decision boundary flexes to account for the training observations (values between 2^-13 and 2^5 with a power-2 transformation).

For NN, there are quite a number of parameters, involving for instance different weight initialization schemes (glorot normal, he normal, glorot uniform etc.), the optimizer (rmsprop, adam etc.) or regularization techniques like dropout, and we only focused on some of them: i) the number of epochs, i.e., of training periods (between 25 and 200), ii) the number of layers before the final layer (between 1 and 3), iii) the number of neurons in the first layer (between 5 and 100), iv) if relevant, the number of neurons in the second layer (between 5 and 100), v) if relevant, the number of neurons in the third layer (between 5 and 100), vi) the drop-out rate for each of the previous layer (one value shared by all layers, between 0.1 and 0.5) and vii) the input drop-out rate for - as the name suggests - the inputs provided to the initial layer (between 0.1 and 0.5)^4^.

For xgboost, we focused on 8 parameters and chose default values for others for the sake of simplicity^5^: i) the maximum number of rounds/iterations, which for classification is similar to the number of trees to grow (between 10 and 1000), ii) the learning rate eta, which shrinks the feature weights after every round, and for which low values slow the learning process down and need to be compensated by an increase in the number of rounds (between 0.01 and 0.2), iii) the regularization parameter gamma, which relies on across-tree information and usually brings improvement with shallow trees (values between 2^-20 and 2^6 with a power-2 transformation), iv) the maximal depth of the tree - the deeper the tree, the more complex problems it can handle, but with a higher risk of overfitting (values between 1 and 10), v) the minimum ‘child weight’, which corresponds to the minimum sum of instance weight of a leaf node (calculated by second order partial derivative), and helps decide when to stop splitting a tree and block potential feature interactions and related overfitting (between 1 and 10), vi) the number of observations supplied to a tree (values between 50% and 100%), vii) the number of features supplied to a tree (values between 50% and 100%), and viii) the regularization parameter alpha, which performs L1 (lasso) regression on leaf weights (values between 2^-20^ and 2^6^ with a power-2 transformation).

### Tuning procedure

There are different methods for exploring a space of hyperparameters. First, a random search can be used, in which a number of acceptable configurations of hyperparameter values, chosen at random, are tried and compared. The number of configurations must increase, however, as the number of hyper- parameters increases and the possible ranges of values widens. In our case, this approach did not appear to be the most efficient. Another option is to perform a grid search, where the range of possible values for each hyperparameter leads to a number of evenly spaced values (possibly with a log/power transformation), and all sets of these values are tried and compared. Again, this method quickly becomes impractical as the number of hyper-parameters increases and one wishes to consider a dense grid. A third option is to perform a tailored search when it makes sense to consider some hyper- parameters first and then others, thus greatly reducing the number of configurations to be explored. Such an approach can be found for instance for xgboost, but not for other techniques. A fourth approach is model-based optimization (MBO), in which a model gradually learns the structure of the hyperparameter space and which hyperparameter values lead to the best performance. This approach is generally much less computationally intensive than previous approaches. A multi-point approach can be considered especially when parallelization of the computations is possible.

We took advantage of a multi-point random-forest-based Bayesian MBO procedure with 25 iterations using a "lower confidence bound" infill criterion, which is suitable for a parameter space that is not purely numerical^6^. For each ‘state-of-the-art’ classifier and each set of predictors, we parallelized the procedure with 25 threads.

Given our tuning approach, we conducted a repeated multiple-fold cross validation with 5 replicates and 5 folds to find the best values of the hyper-parameters. An 80-20 split was adopted for the training and validation sets.

In our case, given our 100 replicates in conjunction with our different algorithms and predictor sets, it was extremely time-consuming to perform the repeated cross-validation procedure each time we divided the whole dataset into a training set and a test set. We therefore decided to separate the hyper- parameter tuning from our assessment of classification performances. For each classifier and each set of predictors, we performed a 5-time repeated 5-fold cross-validation to assess the hyper-parameters. We assume that this provides enough cases, i.e., enough different validation sets, to compute values of the hyper-parameters that perform well on average over many different configurations - our 100 repetitions.

### Performance metrics for tuning the hyper-parameters

A metric was required for the tuning of hyper-parameters. We considered log loss both because it is the only metric in our set of metrics that can be used to train neural networks, and because at a more theoretical level, it corresponds to a ’pure’ Machine Learning perspective, independent from the ’real problem’ (bonobos calls) - log loss as a metric tells us whether the model is confident or not when making a prediction.

As a check, we also used AUC to tune the SVM and xgboost hyper-parameters, and found very similar results to those obtained with log loss.

#### Feature importance

While some algorithms such as xgboost have their own specific techniques for estimating feature importance, we chose a generic method that can be applied to any of our three algorithms SVM, NN and xgboost. It consists in considering each feature in the feature set independently, and for each of them, comparing the quality of the predictions obtained with the initial values with the quality of the predictions obtained after randomly permuting these values across observations. Intuitively, the lower the performance when shuffling, the more important the feature and its values are to classify the observations. This corresponds to the implementation, with the method ‘permutation.importance’, of the function ***generateFeatureImportanceData()*** in the **mlr** package. We chose a number of 50 random permutations to avoid extreme results obtained by chance.

We considered feature importance for both Bioacoustic features and DCT, but left out MFCC due to the number of variables and the difficulty of assigning individual articulatory significance to them.

Again, since we ran 100 iterations for each algorithm and feature set configuration, we averaged the values of feature importance for these interactions to neutralize the sampling error of any given iteration.

#### Ensembling

We considered seven different stacked ensembles:

- For each classifier (SVM, NN and xgboost), we first stacked the models corresponding to the three different feature sets (Bioacoustic parameters, DCT coefficients and MFCC coefficients).
- For each feature set (Bioacoustic, DCT, MFCC), we conversely stacked the models corresponding to the three ‘state-of-the-art’ classifiers.
- Finally, we stacked the 9 models corresponding to the three feature sets processed by the three classifiers.

As for the super learner, we considered a glmnet with alpha = 0 (ridge regression).

#### Evaluating data leakage / BaLi

The Default strategy was implemented through a random sampling procedure which only controlled for equal proportions of the classes of observations in the training and test set. The Fair and Skewed strategies, on the other hand, were implemented with an in-house tool called BALI. Based on the object- oriented R6 framework in R software, BALI relies on a genetic algorithm (with only random mutations and no genome recombination) and the definition of a number of rules - with possibly different weights

- to be respected when creating sets of specified sizes. Some rules are for continuous variables and

aim to equalize or differentiate the mean or variance across rows or columns of different sets. Others deal with categorical variables, and aim to maximize or minimize diversity, etc.

We first defined a rule to ensure similar distributions of occurrences of individuals in the training and test sets. When only this rule is given, the algorithm consistently achieves the objective, with a result similar to the Default strategy. By adding a second rule to prevent or maximize the overlap of sequences on the two sets, we implemented the Fair and Skewed strategies respectively.

#### UMAP and silhouette scores

We used the *umap()* function of the *uwot* package with the default settings, i.e., 15 neighbors, a minimal distance of 0.01, two dimensions and the Euclidean distance metric.

For the silhouette scores (silhouette widths), we used the *get_silhouette()* function of the *clues* package with the Euclidean distance as a measure of dissimilarity between the UMAP coordinates of each data point.

We generated UMAP and silhouette scores for three different configurations: i) the description of the calls with the combined features of the three sets Bioacoustic, DCT and MFCC, ii) the pDFA predictions with the Bioacoustic set, and iii) the predictions of the best ensemble. In the first case, 1,560 inputs were considered - the number of calls in our dataset. In the second and third cases, we considered all the predictions for the calls appearing in the 100 test sets of our approach to compute average performance. For the pDFA, this amounted to 87,500 inputs for call types and 109,000 inputs for individual signatures - i.e., 1,090 and 875 elements in each test set, respectively. For the best ensemble, there were 31,100 inputs for both call types and individual signatures (i.e., 311 elements in each test set, which is the number of elements that were selected by requesting 20% of 1,560 observations with the constraint of identical frequencies of classes in the training and test sets). The rationale behind our approach was that the prediction for a given call could vary across the 100 repetitions, and that keeping this variability was interesting to build a representative and dense space of inputs.

Due to the random nature of the sampling process, the 1,560 calls were not equally represented in the UMAP for pDFA and the best ensemble. On the one hand, when investigating call types, for the pDFA, each call was represented on average 56.09 times, with a minimum of 22 occurrences and a maximum of 85 occurrences. For the best ensemble, each call was represented on average 19.93 times (nearly 20 times, as expected), with a minimum of 9 occurrences and a maximum of 32 occurrences. On the other hand, when assessing individual signatures, for the pDFA, each call was represented 69.87 times on average, with as few as 27 occurrences and as many as 97. For the best ensemble, the average was 19.93 times, the minimum 7 and the maximum 36.

#### Implementation

We relied on the following R packages:

- for the generic ML functions and procedures, ***caret*** (Kuhn, 2008) for pDFA and ***mlr*** (Bischl, et al., 2016) for SV, NN, xgboost and the ensemble learners
- ***MASS*** for the DFA (Venables & Ripley, 2002)
- ***mclust*** for the gmm (Scrucca, et al., 2016)
- ***splitstackshape*** to prepare sets with identical proportions of classes of objects (Mahto, 2019)
- ***mlrMBO*** for the model-based optimization of the hyper-parameters (Bischl, et al., 2017)
- for parallelization of the 100 repetitions to assess performances, ***parallelMap*** (Bischl, Lang, & Schratz, 2021)
- **keras** for the neural networks (CPU-based version 2.8, Python 3.9) (Chollet, et al., 2017)
- ***ggplot2*** for the various figures (Wickham, 2016)
- ***uwot*** for the umap (Melville, et al., 2021) and ***clues*** for the calculation of the silhouette scores (Chang, et al., 2010)
- for data processing broadly speaking, **tidyverse** (Wickham, et al., 2019)
- ***R6*** for the Bali algorithm (Chang, 2022)
- ***missForest*** to impute missing values (Stekhoven, 2022; Stekhoven & Buehlmann, 2012)

All intensive computations – tuning of the hyper-parameters of SVM, NN and xgboost with repeated cross-validation, estimation of the classification performance for simple and stacked learners with 100 runs, estimation of the random baseline with 1,000 runs – were performed on an AMD Ryzen 9 5950X 16-core processor with 32GB RAM running under Windows 11 Pro. 20 threads were used for parallelization during hyperparameter tuning (due to memory constraints), and 25 threads for computing classification performances.

## Acknowledgements

**Funding:** We thank the Ecole Doctorale SIS of the University of Saint-Etienne (Ph.D. grant to SK), the University of Saint-Etienne (research sabbaticals to FL and NM, visiting professorship to VA and research grant), the LABEX ASLAN (ANR-10-LABX-0081) (FL, VA, and FP) of the University of Lyon within the Investments for the Future program (ANR-11-IDEX-0007) operated by the French National Research Agency, the Institut Universitaire de France (NM), and The Canada Graduate Scholarship (XSG) – Michael Smith Foreign Study Supplements (CGS-MSFSS) of the Social Sciences and Humanities Research Council of Canada (SSHRC) and by the Laboratoire Dynamique Du Langage (UMR 5596, University of Lyon 2). We warmly thank the zoological parks of Apenheul, Planckendael and La Vallée des Singes for their welcome, and especially to the bonobo keepers for their support.

## Author Contributions

Conceptualization: all

Data Curation: XSG, FL, SK

Formal analysis: VA, CC, FP, XSG

Funding Acquisition: VA, SK, FL, NM, FP, XSG

Investigation: all

Methodology: all

Project Administration: VA, FL

Software: VA, CC Supervision: VA, CC, FL, FP

Validation: all

Visualization: VA, CC, FL, FP

Writing – Original Draft Preparation: VA, CC, FP

Writing – Review & Editing: VA, CC, FL, NM, FP

## Appendix: Acoustic feature description

**Table A1.**
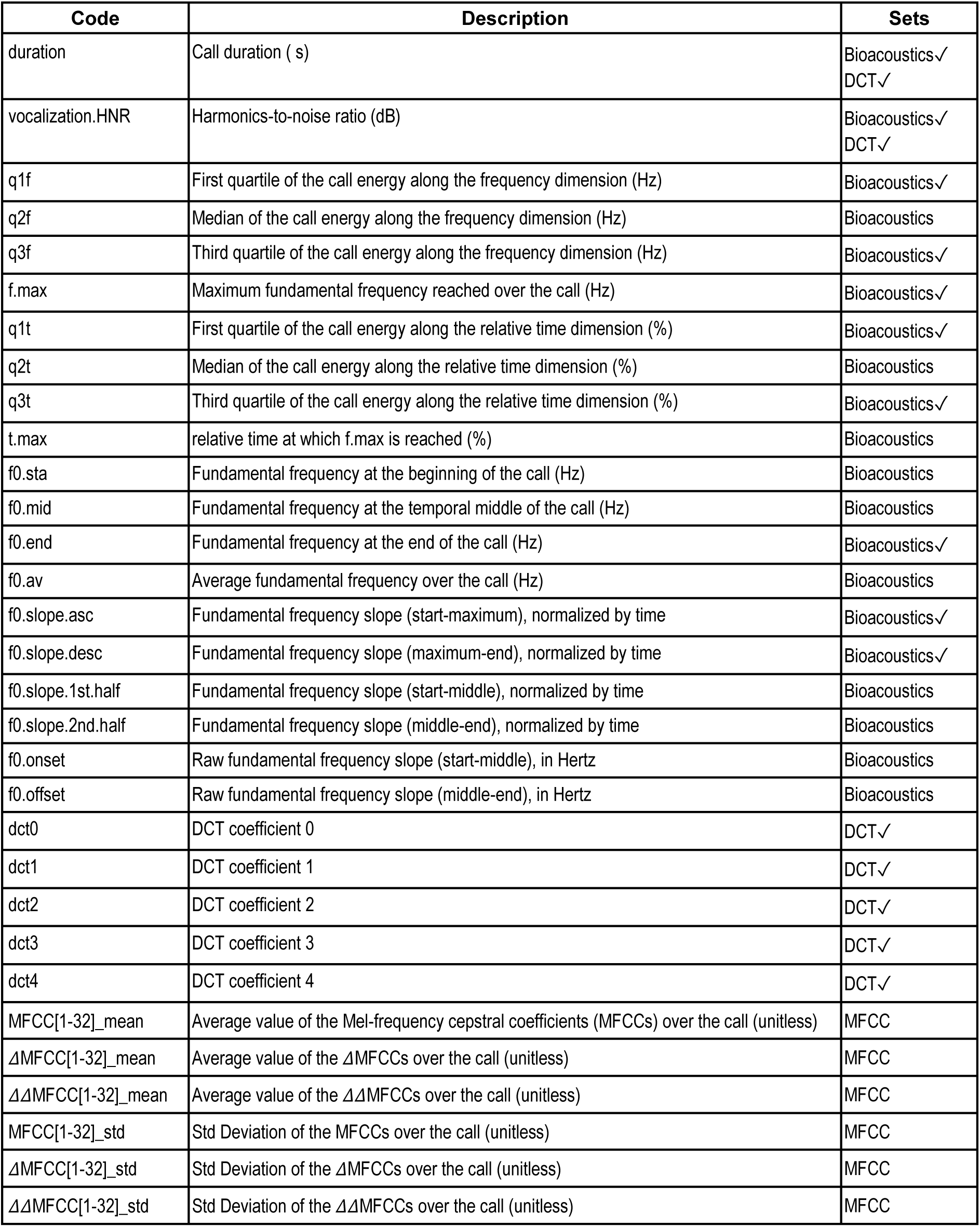
Description of the acoustic features (two first columns) and indication of the feature set composition (third column). A tick indicates that the feature was included in the pDFA analyses (see Methodology pDFA for details).

| Code | Description | Sets |
| --- | --- | --- |
| duration | Call duration ( s) | Bioacoustics✓<br>DCT✓ |
| vocalization.HNR | Harmonics-to-noise ratio (dB) | Bioacoustics✓<br>DCT✓ |
| q1f | First quartile of the call energy along the frequency dimension (Hz) | Bioacoustics✓ |
| q2f | Median of the call energy along the frequency dimension (Hz) | Bioacoustics |
| q3f | Third quartile of the call energy along the frequency dimension (Hz) | Bioacoustics✓ |
| f.max | Maximum fundamental frequency reached over the call (Hz) | Bioacoustics✓ |
| q1t | First quartile of the call energy along the relative time dimension (%) | Bioacoustics✓ |
| q2t | Median of the call energy along the relative time dimension (%) | Bioacoustics |
| q3t | Third quartile of the call energy along the relative time dimension (%) | Bioacoustics✓ |
| t.max | relative time at which f.max is reached (%) | Bioacoustics |
| f0.sta | Fundamental frequency at the beginning of the call (Hz) | Bioacoustics |
| f0.mid | Fundamental frequency at the temporal middle of the call (Hz) | Bioacoustics |
| f0.end | Fundamental frequency at the end of the call (Hz) | Bioacoustics✓ |
| f0.av | Average fundamental frequency over the call (Hz) | Bioacoustics |
| f0.slope.asc | Fundamental frequency slope (start-maximum), normalized by time | Bioacoustics✓ |
| f0.slope.desc | Fundamental frequency slope (maximum-end), normalized by time | Bioacoustics✓ |
| f0.slope.1st.half | Fundamental frequency slope (start-middle), normalized by time | Bioacoustics |
| f0.slope.2nd.half | Fundamental frequency slope (middle-end), normalized by time | Bioacoustics |
| f0.onset | Raw fundamental frequency slope (start-middle), in Hertz | Bioacoustics |
| f0.offset | Raw fundamental frequency slope (middle-end), in Hertz | Bioacoustics |
| dct0 | DCT coefficient 0 | DCT✓ |
| dct1 | DCT coefficient 1 | DCT✓ |
| dct2 | DCT coefficient 2 | DCT✓ |
| dct3 | DCT coefficient 3 | DCT✓ |
| dct4 | DCT coefficient 4 | DCT✓ |
| MFCC[1-32]_mean | Average value of the Mel-frequency cepstral coefficients (MFCCs) over the call (unitless) | MFCC |
| ΔMFCC[1-32]_mean | Average value of the ΔMFCCs over the call (unitless) | MFCC |
| $\Delta\Delta$MFCC[1-32]_mean | Average value of the $\Delta\Delta$MFCCs over the call (unitless) | MFCC |
| MFCC[1-32]_std | Std Deviation of the MFCCs over the call (unitless) | MFCC |
| ΔMFCC[1-32]_std | Std Deviation of the ΔMFCCs over the call (unitless) | MFCC |
| $\Delta\Delta$MFCC[1-32]_std | Std Deviation of the $\Delta\Delta$MFCCs over the call (unitless) | MFCC |

## Footnotes

1 MFCCs were computed with the voicebox toolbox in Matlab, downloaded from https://github.com/ImperialCollegeLondon/sap-voicebox in November 2019. The command v_melcepst(S,Fs,’dD’,32,33,1024,512,500/Fs,12000/Fs) was applied to the audio signal S, sampled at frequency Fs.

2 The stepwise VIF function used in this study has been built up by Marcus W. Beck and was downloaded from: https://gist.github.com/fawda123/4717702

3 This issue is orthogonal to exerting some control over the two sets as what we did to ensure similar representativity of the classes of observations in both train and test the two sets (e.g., when classifying individuals, 23% (362/1560) of the calls should be JIill’s in both sets).

4 To complete the specification of our NN: the final layer contained a number of units equal to the number of classes of the classification problem. A softmax activation was used for this layer in relation to a cross-entropy loss function. We choose a size of 128 for the mini-batches, a glorot uniform weight initialization scheme, and an Adam optimizer with a learning rate of 0.001.

5 Hyper-parameter tuning was the most computationally demanding for xgboost because we chose to work with large ranges of values, which in some extreme cases required a very large number of calculations.

6 We actually initially compared several of the discussed approaches, and found that relying on a model- based optimization led to the best ratio of performances over time.

